# Characterization of neuronal synaptopodin reveals a myosin V-dependent mechanism of synaptopodin clustering at the post-synaptic sites

**DOI:** 10.1101/526509

**Authors:** Judit González-Gallego, Anja Konietzny, Alberto Perez-Alvarez, Julia Bär, Urban Maier, Alexander Drakew, John A. Hammer, Jeroen A. A. Demmers, Dick H. W. Dekkers, Michael Frotscher, Matthias Kneussel, Thomas Oertner, Wolfgang Wagner, Marina Mikhaylova

## Abstract

The spine apparatus (SA) is an endoplasmic reticulum-related organelle which is present in a subset of dendritic spines in cortical and pyramidal neurons. The synaptopodin protein localizes between the stacks of the spine apparatus and is essential for the formation of this unique organelle. Although several studies have demonstrated the significance of the SA and synaptopodin in calcium homeostasis and plasticity of dendritic spines, it is still unclear what factors contribute to its stability at the synapse and whether the SA is locally formed or it is actively delivered to the spines. In this study we show that synaptopodin clusters are stable at their locations. We found no evidence of active microtubule-based transport for synaptopodin. Instead new clusters were emerging in the spines, which we interpret as the SA being assembled on-site. Furthermore, using super-resolution microscopy we show a tight association of synaptopodin with actin filaments. We identify the actin-based motor proteins myosin V and VI as novel interaction partners of synaptopodin and demonstrate that myosin V is important for the formation and/or maintenance of the SA.

**SUMMARY STATEMENT:** Here we demonstrate that the spine apparatus is assembled locally in dendrites. This process, which relies on the presence of synaptopodin and F-actin, is disrupted by interfering with myosin-V activity.

## INTRODUCTION

As an adaptation to their enormous size, neurons have developed a highly sophisticated trafficking system mediating long-distance transport and local control of membrane and protein turnover (Hanus and Ehlers, 2016). Thus, in addition to the somatic localization, most components of the secretory membrane organelles are also found in dendrites (Hanus et al., 2014; Mikhaylova et al., 2016). One key organelle is the endoplasmic reticulum (ER), which in all eukaryotic cells is important for lipid and membrane protein synthesis, as well as protein folding and transport. Additionally, the ER plays a key role in calcium homeostasis. In neurons, the highly complex ER spans the entire cell, including soma, axon and dendritic tree, and is even found in dendritic spines (Toresson and Grant, 2005). Targeting of the ER to spines varies between different neuronal cell types: almost every dendritic spine of cerebellar Purkinje cells contains a tubule of smooth ER associated with the spinous actin cytoskeleton (Wagner et al., 2011a). In contrast, the majority of dendritic spines of pyramidal neurons in the cortex and hippocampus do not contain ER (Toresson and Grant, 2005). However, a subset of spines in cortex and hippocampus contains a complex ER-based organelle called the spine apparatus (SA / Deller et al., 2003). The SA is usually localized in the spine neck or at the base of the spine head and consist of laminar ER stacks with intervening electron-dense plates connected to the main ER network. This intriguing structure serves as a calcium store and is important for synaptic plasticity (Deller et al., 2003). Importantly, the SA can form not only *in vivo* and in organotypic hippocampal slice cultures but also in primary hippocampal neurons (Korkotian et al., 2014).

The protein synaptopodin was identified as an essential component of the SA. It is required for the formation and maintenance of this organelle. Electron microscopy studies in neurons indicated that synaptopodin is localized in between the ER stacks of the SA. It plays an instrumental role in establishing the SA structure, since dendritic spines of synaptopodin-deficient mouse mutants contain single ER tubules but are devoid of the complex SA, and expression of synaptopodin is sufficient to rescue this phenotype (Deller et al., 2003, 2007; Vlachos et al., 2013). Accordingly, cerebellar Purkinje cells, which do not express synaptopodin, do not form SA despite of having ER tubules in all of their numerous spines. Synaptopodin is also involved in the establishment of a similar ER-based structure, called the cisternal organelle, in the axon initial segment (Bas Orth et al., 2007). Synaptopodin is restricted to telencephalic neurons and renal podocytes. Its expression in the brain is developmentally regulated and coincides with synaptic maturation (Czarnecki et al., 2005; Mundel et al., 1997). In humans and rodents, three splice isoforms of synaptopodin have been identified, but only the shorter protein is found in the brain (Asanuma et al., 2005; Chalovich and Schroeter, 2010 / Figure S1). In mouse this isoform corresponds to a 690 aa product (identifier: Q8CC35-3; Figure S1). Synaptopodin is a cytosolic protein. It lacks a clear domain organization, contains several predicted disordered regions, and is known as an actin and α-actinin binding protein (Asanuma et al., 2005; Chalovich and Schroeter, 2010; Kremerskothen et al., 2005). It has been demonstrated that about one third of dendritic spines contains synaptopodin and that it is more frequently present in a subset of spines with a large spine head volume (Vlachos et al., 2009). Long-term imaging of adult hippocampal primary neurons transfected with GFP-synaptopodin revealed that synaptopodin clusters appear to exit or enter dendritic spines during a three-day imaging period (Vlachos et al., 2009), although the imaging interval used in this study did not allow to obtain more detailed trafficking characteristics of synaptopodin. Interestingly, the presence of synaptopodin correlated with synaptic strength, indicating that the SA-positive spines might have different plastic properties (Korkotian et al., 2014; Vlachos et al., 2009). The presence of synaptopodin was shown to regulate the diffusion of metabotropic mGluR5 glutamate receptors in and out of spines (Wang et al., 2016). Along the same line, synaptopodin -/- mice show impaired long-term potentiation (LTP) and spatial learning (Deller et al., 2003; Korkotian et al., 2014).

Notably, synaptopodin mRNA and protein levels are regulated by neuronal activity. For example, it has been shown that in dentate granule cells synaptopodin expression is upregulated following LTP *in vivo* (Yamazaki et al., 2001). Moreover, a recent study has found that synaptopodin is required for cAMP-mediated long-term synaptic potentiation (LTP) in developing neurons and that it is most likely a substrate of protein kinase A (PKA), which becomes activated during LTP (Zhang et al., 2013). In renal podocytes, it was shown that synaptopodin can be phosphorylated by protein kinase A (PKA) and CaMKII. Together with the phosphatase calcineurin, these kinases regulate the phosphorylation status of synaptopodin and its association with the protein 14-3-3, which can protect synaptopodin from degradation via cathepsin L (Faul et al., 2008). Therefore, it is plausible that, in active dendritic spines, mechanisms similar to those in podocytes regulate synaptopodin stability and protect it from degradation. Taken together, the data suggest that synaptopodin acts as a powerful tool to induce formation of the SA in dendritic spines and it is very likely that synaptopodin expression, localization, function, and stability are highly regulated.

Overall, despite the importance of the SA in synaptic function, there are still many open questions about the origin of this organelle. For instance, is the complete SA actively transported along the dendrite and then targeted to selected spines, or it is assembled locally as needed? And what are the molecular mechanisms that regulate SA localization?

In this study we aimed to address these questions and to learn more about the dynamics of the spinous ER and the SA in hippocampal neurons. 2-photon imaging in hippocampal slice cultures indicated that the majority of spines containing ER were also positive for synaptopodin. Next, we performed long-term live-cell imaging of primary neurons transfected with GFP-synaptopodin and to our surprise found no evidence of synaptopodin clusters being actively transported along dendritic branches. Simultaneous live imaging of the ER and synaptopodin revealed a gradual formation of synaptopodin clusters in spines, which we interpret as the SA being assembled on-site. Using super-resolution microscopy, we show that synaptopodin in dendrites and spines is often embedded in dense F-actin mesh. In order to identify factors that allow for synaptopodin localization at postsynaptic sites, we performed a mass-spectrometric analysis of brain-specific binding partners isolated via a pull-down assay. Interestingly, several myosins were found in the analysis, including the processive motors myosin V and VI. Using dominant-negative approaches, pharmacology and knock out mice, we show that myosin VI is dispensable for the spine localization of synaptopodin, whereas myosin V affected the formation and/or maintenance of synaptopodin clusters resulting in diminished synaptic targeting of synaptopodin.

## RESULTS

### Synaptopodin is mainly present in spines containing ER and is upregulated during neuronal development

In organotypic hippocampal slice cultures, only synapses on spines containing ER undergo long-term depression mediated by activation of type 1 metabotropic glutamate receptors, which is accompanied by calcium release events of large amplitude (Holbro et al., 2009). Here we ask how the presence of ER in dendritic spines correlates with the presence of synaptopodin. 2-photon imaging of hippocampal CA1 neurons electroporated with plasmids encoding tdimer2 as a morphology marker and ER-GFP as a label for the ER followed by post-hoc fixation and immunolabeling with synaptopodin-specific antibody indicated that 71% of spines containing ER were also synaptopodin positive and only 5% of spines without ER contained synaptopodin (Figure 1A-C). Synaptopodin labeling is frequently used as a marker for the SA (Deller et al., 2000, 2003), therefore we conclude that these spines contain a SA.

**Figure 1.**
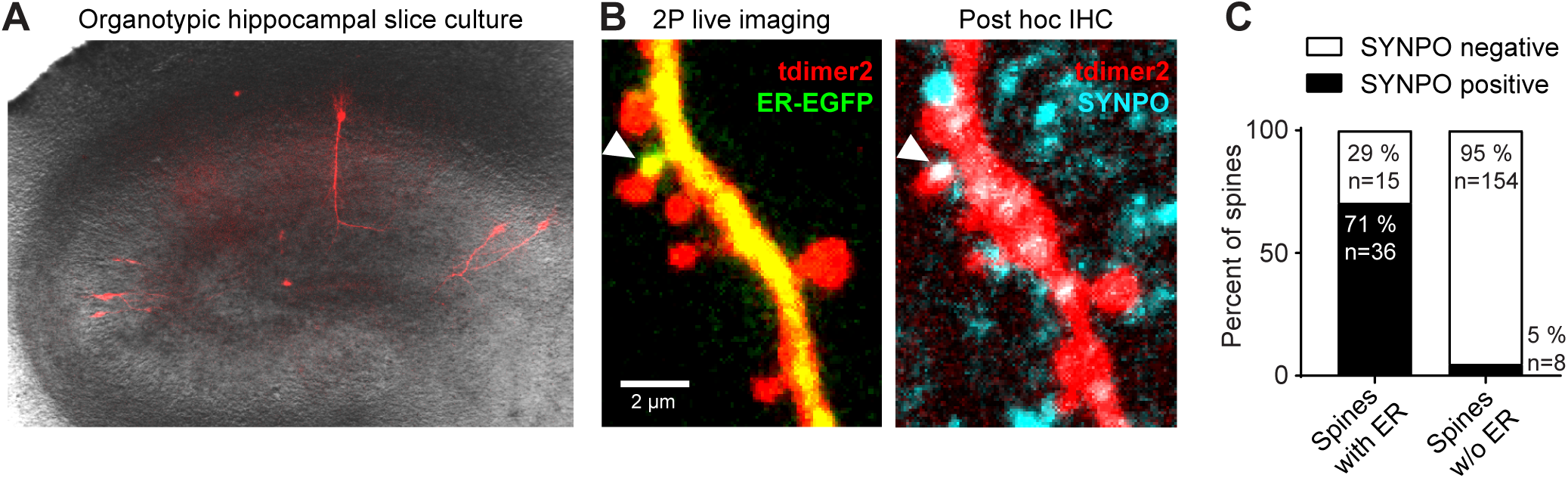
Synaptopodin is mainly present in spines with endoplasmic reticulum (ER) (A) Overlay of DIC and epifluorescence image of an organotypic hippocampal slice with few CA1 pyramidal neurons expressing tdimer2 and ER-EGFP. (B) Two-photon live imaging from a CA1 neuron apical dendritic segment (left) and post hoc antibody staining against synaptopodin (right). Arrowhead shows a spine containing ER with positive staining for synaptopodin. Scale bar = 2 µm. (C) Quantification of synaptopodin staining in spines according to presence or absence of ER in 2P live imaging (2 slices, 2 neurons).

The expression of synaptopodin is developmentally regulated and coincides with synaptogenesis. It has been reported that in rat hippocampal primary neurons, synaptopodin is first found at day *in vitro* (DIV) 12 and the expression reaches its maximum after DIV25, however the authors did not show the data (Mundel et al., 1997). In another study synaptopodin was detected in dendritic spines and the shaft of DIV10 hippocampal neurons (Vlachos et al., 2009). In both cases, the cultures were prepared using classical neuronal culturing medium supplemented with B27. Recently a new formulation of a serum-free synthetic supplement, SM1, for primary neuronal cultures was introduced (StemCell Technologies). SM1 promotes synaptic development and neuronal activity and is better suited for studying synaptic function *in vitro*. Therefore, we decided to establish a time-line of synaptopodin expression in hippocampal neurons prepared from E18 rat embryos and grown in medium supplemented with SM1. Confocal imaging showed that at DIV10, a clear synaptopodin signal was detectable in dendrites labeled by MAP2 (Figure 2A, Figure S2). The total number of puncta increased until DIV15. At DIV21 we did not see more synaptopodin puncta than at DIV15, but instead more puncta were found in areas outside of the dendritic shaft, which could suggest an increased localization to dendritic spines (Figure 2B-C). Therefore, we next included the postsynaptic marker homer1 and the F-actin marker phalloidin, to visualize dendritic spines. We observed a trend to increased colocalization of synaptopodin with homer1 in mature cultures (Figure 2D-E). On average about 48% of synaptopodin puncta were adjacent to homer1-positive synapses in DIV10 neurons, while this number increased to 64% and 62% at DIV15 and DIV21, respectively (Figure 2E), indicating an increase in synaptic association of synaptopodin clusters. Interestingly, we also observed a large number of synaptopodin clusters colocalizing with homer1 in locations that seem to be inside the dendritic shaft, as opposed to spine localization (Figure 2D-E). Some of those certainly can be accounted for by spines that reach out orthogonally to the plane of view. However, such spines are generally rare in primary neuronal culture, and we speculate that some of these sites might constitute excitatory shaft synapses.

**Figure 2.**
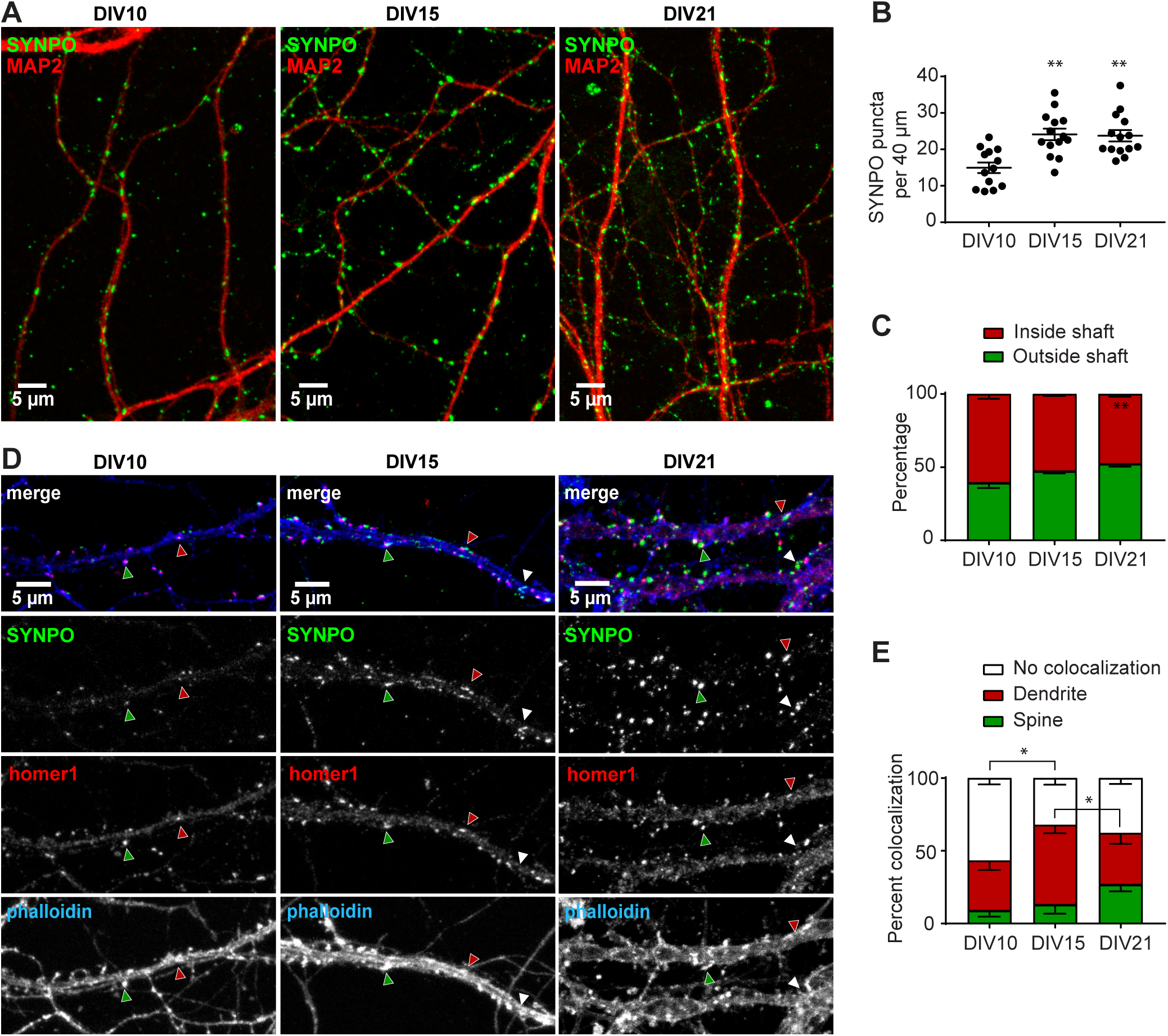
Expression and localization of synaptopodin during development in hippocampal primary neurons. (A) Representative confocal images of primary hippocampal neurons on DIV10, DIV15 and DIV21 stained with anti-synaptopodin and anti-MAP2 antibodies. Scale bar = 5 µm. (B) Quantification (mean ± SEM) of the average number of total synaptopodin puncta per 40 µm dendritic segments at DIV10, DIV15 and DIV21. Kruskal-Wallis-Test, p=0.0005, ** p=0.0012 (DIV10 vs DIV15), p=0.0035 (DIV10 vs DIV21). (C) Quantification (mean ± SEM) of synaptopodin puncta present inside or outside (spines, filopodia) of dendritic shafts. Kruskal-Wallis-Test, ** p=0.0062 (DIV10 vs DIV21). (B+C) DIV10: n = 14 cells from 3 independent cultures with 63 separate segments counted. DIV15: n = 14 cells from 2 independent cultures with 93 separate segments counted. DIV21: n = 16 cells from 3 independent cultures with 100 separate segments counted. (D) Representative confocal image of primary hippocampal neurons at DIV10, DIV15 and DIV21 stained with anti-synaptopodin, anti-Homer1 and phalloidin-A647N. Red arrows: colocalization in spine. Green arrow: colocalization in dendrite. White arrow: no colocalization. Scale bar = 5 µm. (E) Quantification (mean ± SEM) of the percentage of synaptopodin puncta colocalizing with Homer1 in different dendritic subcompartments. Mixed ANOVA with DIV as between and localization as within group factors. F(4, 36)=4.4359, p=0.00512. Newman-Keuls post hoc test: * p=0.0348 (no coloc DIV10 vs DIV15), p=0.0435 (DIV15 vs DIV21 dendrite). DIV10: n = 8 cells from 3 independent cultures with 44 separate segments counted. DIV15: n = 7 cells from 2 independent cultures with 35 separate segments counted. DIV21: n = 8 cells from 2 independent cultures with 41 separate segments counted.

These data indicate that similarly to the previous reports, synaptopodin expression is increasing during synaptogenesis and synaptic maturation in primary neuronal culture, and DIV15-21 neurons can be used as a cell culture model to study mechanisms regulating synaptopodin cluster localization and formation in dendritic spines.

### Synaptopodin clusters show no processive trafficking and are stably anchored on the spinous and dendritic F-actin

As shown previously, the distribution of transfected GFP-synaptopodin closely matches the distribution of the endogenous protein (Vlachos et al., 2009). Here we generated a construct for GFP-synaptopodin expression under the human synapsin 1 promoter, which is frequently used for low-to-moderate neuron-specific expression of proteins in primary neurons and organotypic slices (Kügler et al., 2003; Mikhaylova et al., 2016). We co-transfected DIV14 hippocampal neurons with GFP-synaptopodin and mRuby2 as a morphology marker and fixed the cells 24 hours later. Confocal imaging indicated that, in agreement with endogenous synaptopodin localization, GFP-synaptopodin clusters were present in a subset of dendritic spines, sometimes formed clusters in dendritic shafts and, as reported earlier for the endogenous protein, are also present at the axonal initial segment (Figure S3; Bas Orth et al., 2007; Sánchez-Ponce et al., 2011).

Next, we asked whether GFP-synaptopodin clusters are actively transported within the dendrite, and how they are recruited into or removed from dendritic spines. Surprisingly, continued time-lapse imaging of GFP-synaptopodin and mRuby2 co-transfected neurons did not reveal a single long-range transport event in dendrites over the 7 hours of imaging with a 5 min interval (Figure 3A). Synaptopodin puncta in both spines and dendrite shafts were stably anchored at the same locations and sometimes oscillated within areas of 1-2 µm. These data rule out the possibility that clusters of synaptopodin (and thus the SA) might be actively transported via long-distance microtubule-dependent, active transport as it is known for many other membrane organelles (Ayloo et al., 2017; Valenzuela and Perez, 2015; van Spronsen et al., 2013).

**Figure 3.**
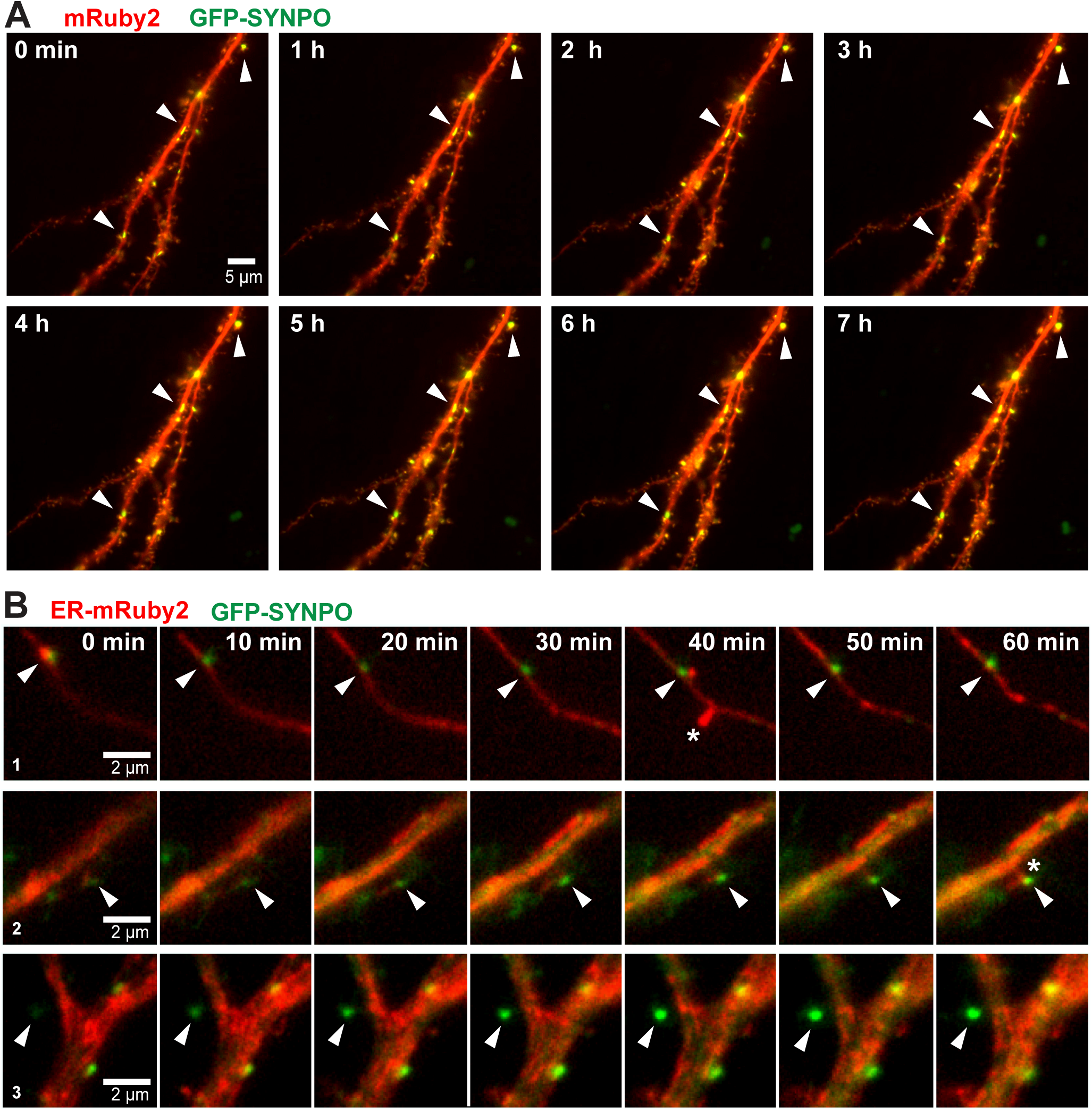
Synaptopodin puncta are stably anchored over prolonged time periods and generated *de novo* in dendritic spines. (A) Time-lapse imaging of a primary hippocampal neuron (DIV15) transfected with mRuby2 and GFP-synaptopodin followed over 7 hours with 5 min intervals. Shown are frames 1 h apart, no changes were observed between these timepoints. Arrowheads indicate examples of GFP-synaptopodin puncta that were stable over the entire imaging period. Scale bar = 5 µm. Also see Movie 1. (B) Time-lapse imaging of a primary hippocampal neuron (DIV15) expressing GFP-synaptopodin and an ER marker (ER-mRuby2). 1: Arrowhead: Synaptopodin colocalizes with spinous ER protrusion. Asterisk: ER transiently entering spine. 2: Synaptopodin localized in spine that experiences ER entry (asterisk). 3: Intensity of spine-localized synaptopodin increases over time. Scale bar = 2 µm. Also see Movie 2.

This leaves the scenario in which the SA is formed locally *de novo*, potentially via the recruitment of synaptopodin, in a subset of spines visited by the ER. To test this, we co-transfected primary neurons with the ER marker fused to mRuby2 and GFP-synaptopodin. Time-lapse imaging showed that similarly to organotypic slices (Figure 1), ER tubules were entering protrusions coming from the dendritic shaft, most likely spines or filopodia (Figure 3B, example 1, Movie S2). In some cases, ER entry into a spine coincided with an increase of synaptopodin intensity in that spine (Figure 3B, example 2, Movie S2). Alternatively, in rare cases the synaptopodin signal gradually emerged in a spine without detectable ER (Figure 3B, example 3, Movie S2). In analogy with organotypic slices, this could be dendritic spines with synaptopodin which do not contain the ER (Figure 1). However, since the images were taken with 5 min intervals we cannot completely rule out that this spine was visited by the ER between the acquisition frames (Movie S2). Taken together, it is very likely that synaptopodin and the ER are locally emerging at synaptic sites to form a SA. Of note, also in this experiment we found no evidence of large synaptopodin clusters being transported.

Synaptopodin is an actin-associated protein (Asanuma et al., 2005; Kremerskothen et al., 2005; Mundel et al., 1997). However, the detailed spatial relation of synaptopodin puncta to neuronal F-actin could not be directly resolved due to the diffraction limit of fluorescence imaging techniques used in earlier studies (Verbich et al., 2016; Vlachos et al., 2009). Super-resolution imaging based on stochastic optical reconstruction microscopy (STORM) and photoactivated localization microscopy (PALM) were recently used to show the association of overexpressed Dendrasynaptopodin with F-actin in a neck of dendritic spines in hippocampal primary neurons (Wang et al., 2016). Here we decided to visualize endogenous synaptopodin by employing simulated emission depletion (STED) nanoscopy. To this end, we stained hippocampal primary neurons with an antibody against MAP2 and synaptopodin, while the F-actin cytoskeleton was labeled by phalloidin-Atto647N. Two-color STED imaging of synaptopodin and F-actin indicated that synaptopodin-labeled structures were tightly embedded into the F-actin mesh irrespectively of spinous or shaft localization (Figure 4A-B). This association with actin filaments within spines and dendritic shafts suggests that the immobility of synaptopodin puncta may be due to anchoring at the actin cytoskeleton.

**Figure 4.**
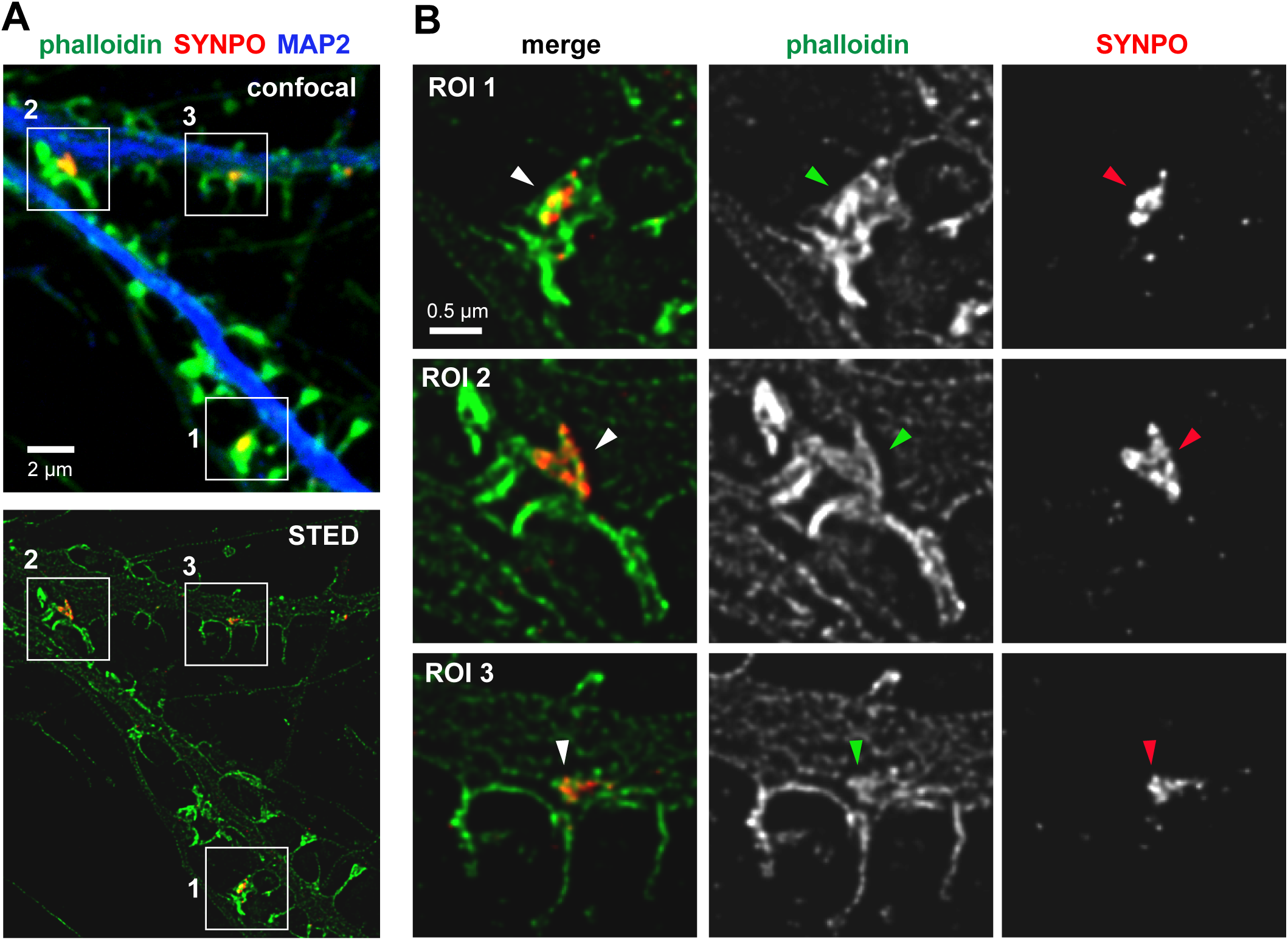
Synaptopodin exists in close association with F-actin in spines and inside dendritic shafts. (A) Confocal and corresponding STED images of DIV42 primary hippocampal neuron with F-actin stained by Phalloidin-A647N and antibody-labelling of endogenous synaptopodin. White boxes indicate the positions of enlarged insets in (B). 1: synaptopodin with F-actin inside spine head. 2+3: synaptopodin with F-actin at the dendritic base of a spine. ROI = region of interest. Scale bar = 2 µm (A), 0.5 µm (B).

### Processive myosins are novel interactors of synaptopodin

Anchoring of synaptopodin at actin filaments within spines and dendritic shafts suggests that actin binding proteins might be involved in targeting the SA to specific localizations (Kremerskothen et al., 2005; Mundel et al., 1997). Previously, an interaction between the long splice isoform of human synaptopodin (identifier: Q8N3V7-2; 903 aa) and α-actinin-2 has been suggested to mediate localization of synaptopodin at spines via a binding motif localized between aa 752-903 (Kremerskothen et al., 2004). A truncated version (aa 483 – 752) no longer showed binding to α-actinin and lost dendritic spine localization. However, in our hands, and as reported in several other studies, the short splice isoform of mouse synaptopodin (identifier: Q8CC35-3; 690 aa), lacking the proposed targeting domain of the truncated version, is clearly enriched at spines (Figure S1, 3; Korkotian et al., 2014; Vlachos et al., 2009). Furthermore, Asanuma et al. showed that the short splice isoform contains two α-actinin-2 and α-actinin-4 binding sites of its own (Asanuma et al., 2005). Thus, whether α-actinin plays a direct role in localizing the SA to selected spines still remains to be shown conclusively. However, α-actinin is enriched in all types of spines, whereas synaptopodin is present only in a subset (Hodges et al., 2014; Matt et al., 2018; Nakagawa et al., 2004). Thus, the question of synaptopodin targeting remains open.

In order to obtain insight into the molecular mechanisms that govern the distribution of synaptopodin, we set out to obtain unbiased information about the brain-specific synaptopodin interactome (690 aa isoform). As a first step, we performed a mass spec pull down from rat hippocampus using biotinylated synaptopodin produced in HEK293 cells (Figure 5A-B). We found several interaction partners that have been reported previously to bind synaptopodin, including actinins, actin and 14-3-3 eta (Table S1; Table S2; Figure 5C). Interestingly, CamKIIα and CamKIIβ isoforms were also present in complex with synaptopodin. In renal podocytes, synaptopodin is known as a substrate of CamKII (Faul et al., 2008). Our data indicate that this interaction might also take place in neurons. We found that several actin stabilizing, capping, severing and modifying proteins including tropomodulins (Tmod2, Tmod3), gelsolin (Gsn), Arp2/3 complex (Arpc2), coronins (Coro2a, Coro2b), F-actin-capping proteins (Capza1, Capza2) were specifically enriched in the synaptopodin pull-down (Figure 5C, Table S1, Table S2). This finding indicates that the association of synaptopodin and actin filaments might be more complex than just direct binding to actin and can be subjected to regulation. On the other hand, there were almost no synaptic scaffolding molecules, except Sipa1l1 present in this fraction. In line with our life imaging experiments, we found no kinesin or dynein motor proteins or their adapters present in the synaptopodin complex (Table S1). This might explain why we did not see processive long-distance trafficking of synaptopodin clusters. Out of the novel prominent binding partners, particularly unconventional myosins attracted our attention (Figure 5C). It has been shown that myosin Va is required for targeting of the ER into the spines of cerebellar Purkinje neurons (Wagner et al., 2011a). We hypothesized that the processive myosins V and VI also transport and anchor synaptopodin to dendritic spines thereby mediating the synaptopodin-dependent formation of the spine apparatus. To verify these interactions, we performed the pull-down assay from rat hippocampus, and analyzed individual interactors via western blot. This confirmed the association of bio-GFP-synaptopodin but not the bio-GFP control with the endogenous myosin Va, Vb, VI and Id (Figure 5D). Processive myosin V and VI are highly expressed in pyramidal neurons. They mediate organelle trafficking in and out of dendritic spines as well as provide stable anchoring on actin filaments (Correia et al., 2008; Esteves da Silva et al., 2015). Therefore, we decided to investigate the role of these myosins in the synaptic targeting of synaptopodin.

**Figure 5.**
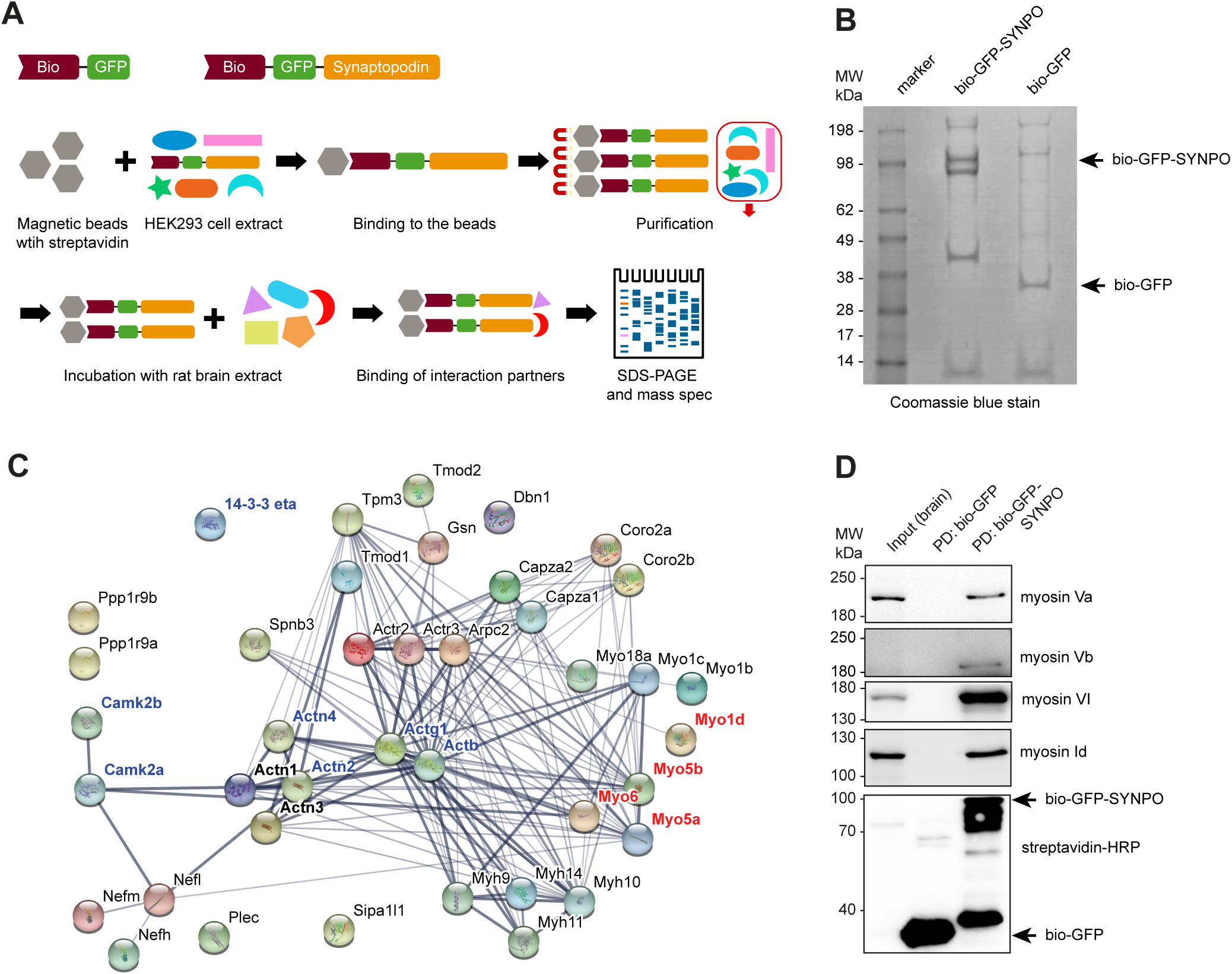
Mass-spectrometry analysis of brain-specific synaptopodin binding partners. (A) Pull-down workflow. Biotinylated GFP-synaptopodin was expressed in HEK cells and bound to Streptavidin-beads. The beads were then incubated with rat brain lysate to pull down brain-specific interactors of synaptopodin. (B) SDS-poly-acrylamide gel of the bio-GFP-synaptopodin and bio-GFP (control) pull-down used for mass-spectrometric analysis. (C) Network analysis of selected potential synaptopodin-interactors identified in the mass spectrometry analysis using the online STRING analysis tool. Highlighted in blue are known synaptopodin-interacting proteins including actin (Actg1, Actb) and actinins (Actn2, Actn4). In red are myosins that were later tested positively in the co-immunoprecipitation in (D). Line thickness indicates strength of data support. (D) Western blot analysis of bio-GFP-synaptopodin pull-down from brain lysate (as in (A)) confirms interaction with myosin Va, Vb, VI and Id. Input = brain lysate, Strept-HRP = streptavidin coupled to horse-radish peroxidase. Arrows indicate expected MW of bio-GFP-synaptopodin (upper) and bio-GFP (lower).

### Synaptopodin cluster numbers and their short-range motility depend on myosin V

To test the contribution of myosin V and VI to the localization of synaptopodin clusters we used a dominant-negative (DN) approach that is well established to inhibit these myosins’ functions (Osterweil et al., 2005; Wu et al., 2002). The DN effect is achieved through overexpression of the C-terminal myosin cargo binding domain, which likely competes with the endogenous, functional myosin for cargo binding. We transfected DIV15 primary hippocampal neurons with a construct containing the cargo binding domain of myosin VI (MyoVI DN) fused to GFP under a human synapsin 1 promoter. Neurons were fixed 24 h after transfection and stained against MAP2. MyoVI DN showed a diffuse distribution throughout the cell and thus was used as a volume marker (Figure 6A). Although quantification showed a reduction of synaptopodin puncta following overexpression of MyoVI DN, the difference to control neurons was not statistically significant (Figure 6B). Additionally, the distribution of synaptopodin puncta between dendrite shafts and spines did not differ from the mRuby2 transfected control neurons (Figure 6C). To corroborate these findings, we utilized primary hippocampal cultures from wild-type and myosin VI-deficient *Snell’s waltzer* (*Myo6*^*sv/sv*^) mice (Avraham et al., 1995). Neurons were fixed at DIV8, 14 and 17 and stained against synaptopodin, homer1 and F-actin (Figure 6D). Quantification of confocal images showed that the number of synaptopodin puncta increased during development in both wild-type and *Myo6*^*sv/sv*^ dendrites, and that the numbers do not significantly differ between the genotypes (Figure 6E). Also the distribution of homer1-colocalized synaptopodin clusters between spines and dendritic shafts was not different in *Myo6*^*sv/sv*^ neurons compared to control (Figure 6F). Thus, we conclude that the interaction of synaptopodin with myosin VI is not essential for the formation of synaptopodin clusters or for their normal distribution to dendritic spines.

**Figure 6.**
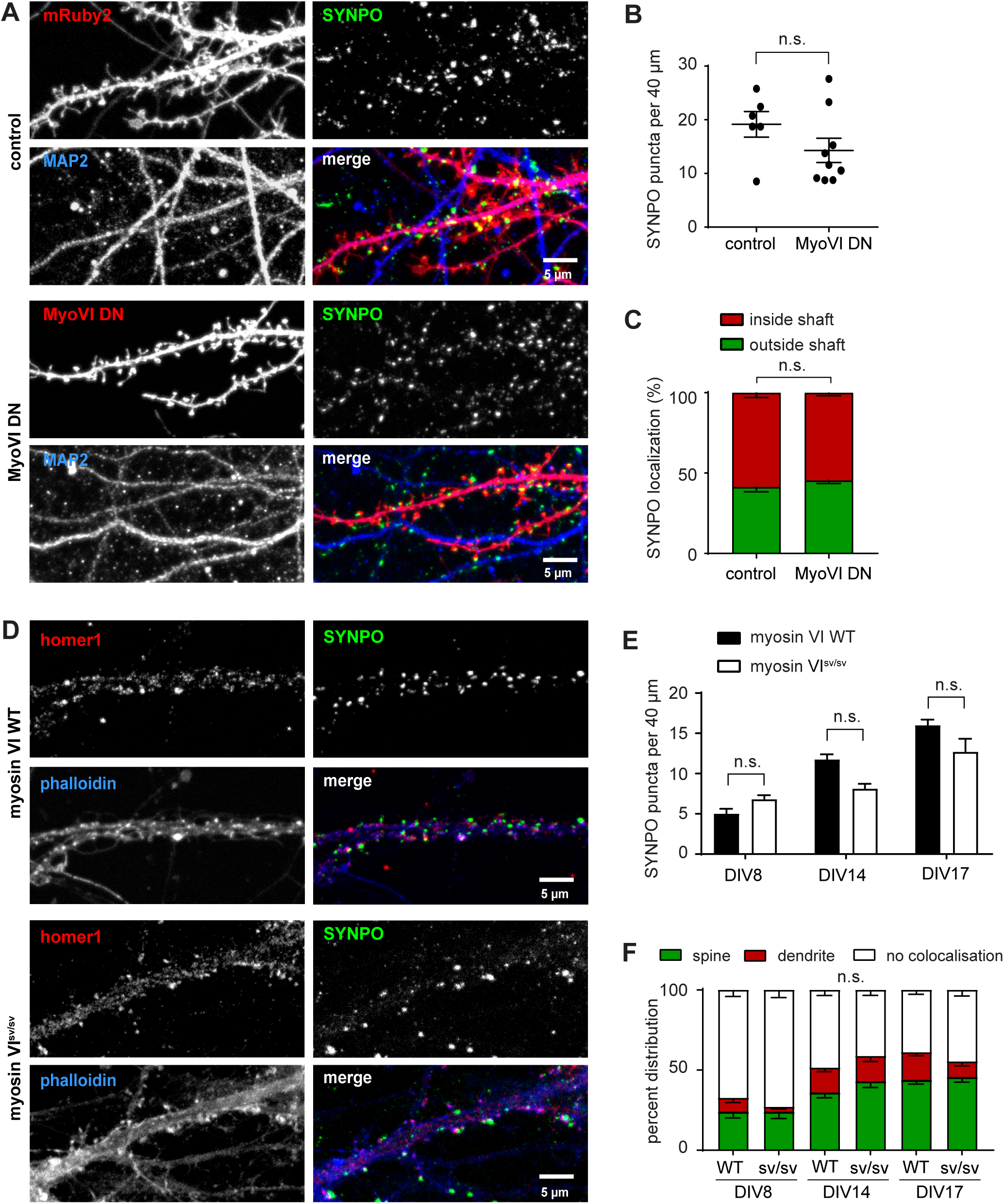
Myosin VI activity is not required for correct localization of synaptopodin. (A) Representative confocal image of primary hippocampal neurons on DIV16 transfected with mRuby2 (control) or GFP-Myosin-VI-dominant-negative (MyoVI DN) and stained with anti-synaptopodin and anti-MAP2. Right column shows single color channels. Scale bar = 5 µm. (B) Quantification (mean ± SEM) of the average number of total synaptopodin puncta per 40 µm dendritic segments in control and MyoVI DN-expressing neurons. Mann-Whitney test, p=0.3277 (n.s.). (C) Quantification of synaptopodin puncta present inside of dendritic shafts or outside (spines, protrusions). Unpaired t-test (two-tailed), p=0.2370. (B+C) MyoVI DN: n=9 cells from two independent cultures with 45 segments counted. mRuby: n=6 cells from two independent cultures with 22 segments counted. (D) Representative confocal image of primary hippocampal neurons on DIV17 from wild type (WT) and myosin VI-deficient (*Myo6*^*sv/sv*^) mice stained with anti-synaptopodin, anti-homer1 and phalloidin-A647N. Right column shows single color channels. Scale bar = 5 µm. (E) Quantification (mean ± SEM) of the average number of total synaptopodin puncta per 40 µm dendritic segments in WT and *Myo6*^*sv/sv*^ neurons. Two-Way ANOVA, wt vs ko p=0.1364 (n.s.) (F) Quantification (mean ± SEM) of the percentage of synaptopodin puncta colocalizing with homer1 inside dendritic shafts or spines in WT and *Myo6*^*sv/sv*^ cultures. Mixed ANOVA with DIV as between and localization as within group factors shows no significant differences. F(2, 22)=1.3869, p=0.271. (E+F) All data comes from two independent cultures. DIV8 WT: n=7 cells with 30 segments counted. DIV14 WT: n=8 cells with 32 segments counted. DIV17 WT: n=8 cells with 47 segments counted. DIV8 KO: n=6 cells with 23 segments counted. DIV14 KO: n=6 cells with 31 segments counted. DIV17 KO: n=8 cells with 28 segments counted.

To examine the functional role of the myosin V-synaptopodin interaction, we again first used a DN approach. Overexpression of the C-terminal myosin Va cargo binding domain, dimerized by the myosin’s coiled coil region (CC), is well known to impair the function of endogenous myosin Va (Balasanyan and Arnold, 2014; Correia et al., 2008; Jones et al., 2000). Here, we developed a novel DN construct comprising only the globular tail domain of myosin Va dimerized by a leucine zipper (LZ) in order to avoid additional effects mediated by the CC region that sequesters further interaction partners from their physiological targets. Since myosin Va is already known to mediate the targeting of ER into the dendritic spines of Purkinje cells, we used this cell type to test for a DN effect (Figure S4). Confocal live imaging of primary mouse Purkinje cells showed that both, constructs with either the myosin Va CC or the LZ fused to the myosin’s globular tail domain, exert a DN effect on ER targeting to spines (Figure S4). Therefore, we decided to use the minimized LZ-globular domain fusion construct (hereafter referred to as MyoV DN) under the human synapsin 1 promoter in hippocampal neurons. MyoV DN sometimes was more enriched at specific dendritic locations, presumably bound to a cargo, but overall entirely filled dendrites and spines and thus could be used as a volume marker (Figure 7A). 24 hours of MyoV DN expression resulted in more diffuse synaptopodin staining and in a significant reduction of synaptopodin clusters compared to control (on average 13.5 per 40 µm in MyoV DN and 24 per 40 µm in mCerulean control; Figure 7B). Importantly, the total number of homer1-positive dendritic spines did not change (Figure 7C). This indicates that myosin Va is required for the formation or stability of synaptopodin clusters and most likely the associated SA.

**Figure 7.**
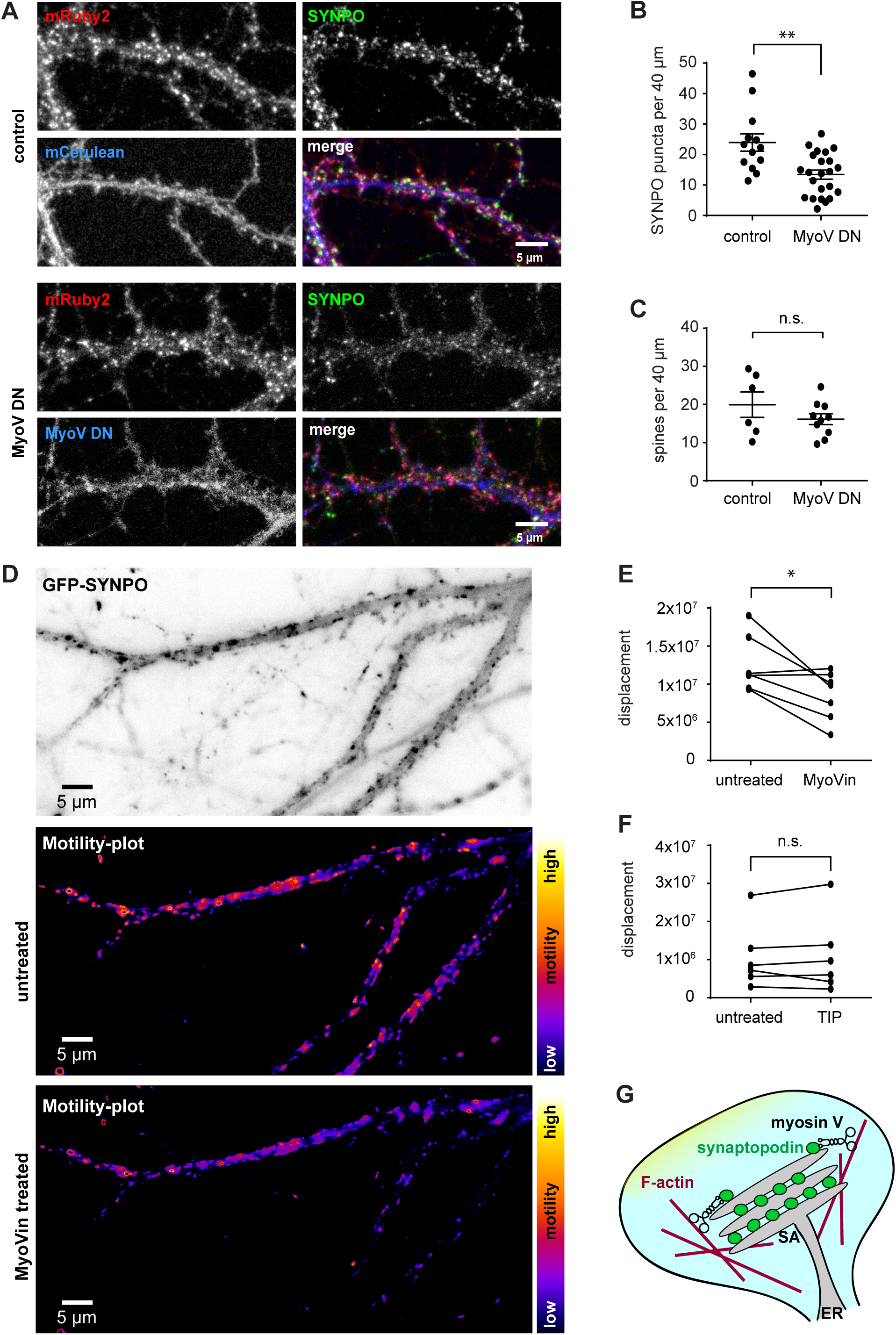
Myosin V inhibition affects quantity and motility of synaptopodin puncta. (A) Representative confocal image of primary hippocampal neurons on DIV16 transfected with mCerulean (control) or Myosin-V-DN-mCerulean (MyoV DN) and stained with anti-synaptopodin and anti-homer. Right column shows single color channels. Scale bar = 5 µm. (B) Quantification (mean ± SEM) of the average number of total synaptopodin puncta per 40 µm dendritic segments in control and MyoV DN-expressing neurons shows a decrease of synaptopodin puncta. Mann-Whitney test, ** p=0.0025. MyoV DN: n=24 cells from 3 independent experiments with 39 segments counted. mCerulean: n=13 cells from 2 independent experiments with 36 segments counted. (C) Quantification (mean ± SEM) of the total number of homer1-positive spines per 40 µm dendritic segments in control and MyoV DN-expressing neurons shows no significant change. Mann-Whitney test, p= 0.4923 (n.s.). One experiment. Control: n = 6 cells with 16 segments counted. MyoV DN: n = 10 cells with 19 segments counted. (D) Representative TIRF image from a timelapse series of a DIV16 primary hippocampal neuron expressing GFP-synaptopodin (top). Motility of synaptopodin-puncta within the same cell before and after 30 min MyoVin treatment (30 µM) was assessed. The motility plots sum up the differences in grayscale values of each pixel from frame to frame. Scale bar = 5 µm (bottom). (E) Quantification (mean ± SEM) of the displacement (shown in D) as integrated density of the motility plot before (untreated) and after treatment. MyoVin treatment reduced the overall motility of synaptopodin puncta. Paired t-test (two-tailed), * p=0.0239. (F) The same experiment as in (D) and (E) was repeated with the myosin VI inhibitor 2,4,6-triiodophenol (TIP, 4 µM) instead of MyoVin. TIP treatment had no effect on the overall motility of synaptopodin puncta. Paired t-test (two-tailed), p=0.7208 (n.s.). (G) Model summarizing the role of myosin V and synaptopodin interaction in localization of the SA to actin filaments associated with the synapse.

Our live imaging experiments have shown that GFP-synaptopodin clusters do not traffic over long distances but that they oscillate in an area of a few micrometers. This type of short-range movements might be mediated by myosin Va, relocating the SA within the spine neck or inside the dendritic shaft as a response to neuronal activity (Vlachos et al., 2009). To test whether these oscillations depend on myosin V we used a pharmacological myosin V inhibitor, MyoVin, which blocks the ATPase activity of the motor domains and can thus turn myosin V from a processive motor into a tether (Gramlich & Klyachko, 2017; Heissler et al, 2017). To increase the number of targeted cells within the same treatment condition we infected hippocampal primary neurons with adeno-associated viruses expressing GFP-synaptopodin and mRuby2 as a cell fill. Imaging of DIV16 neurons for 5 min with 10 sec intervals showed short-range movements of synaptopodin puncta. We then acutely treated the neurons with MyoVin and imaged the same cells 45-60 min after (Figure 7D). Motility analysis indicated that the displacement was significantly reduced following MyoVin application, which is consistent with myosin Va switching from an F-actin motor to a tether (Figure 7E). In line with our previous experiments, the same assay performed with the myosin VI inhibitor 2,4,6-triiodophenol (TIP) showed that TIP has no effect on the displacement of synaptopodin (Figure 7F).

Taken together, this indicates an involvement of myosin V, rather than myosin VI, in the formation or stabilization of synaptopodin clusters and/or the SA, and possibly targeting to specific subcellular sites.

## DISCUSSION

Functional and structural properties of dendritic spines containing the SA differ from their neighbors without the SA (Korkotian et al., 2014; Vlachos et al., 2009). Synaptopodin is closely associated with ER-derived membranes that are forming the SA. It plays a key role in synaptic calcium homeostasis by regulating ER calcium release via ryanodine and inositol-triphosphate receptors (Vlachos et al., 2009). Despite a growing number of studies addressing the role of the SA in neuronal function, it was still enigmatic how the SA gets localized to dendritic spines. Long-term imaging experiments in primary hippocampal neurons revealed that synaptopodin can enter, exit or relocate within the spine over large time spans. In some instances, it seemed that synaptopodin puncta that were initially detected in a spine head had exited the spine and were later found in the shaft (Vlachos et al., 2009.). However, since these data were acquired with a one-day interval, it is difficult to judge whether synaptopodin clusters were actively relocated as a whole, or disassembled and newly assembled elsewhere. In this study we aimed to bridge this gap and learn more about transport and synaptic targeting of synaptopodin.

Imaging of dendritic spines from CA1 hippocampal neurons in slice cultures indicated that the majority of dendritic spines containing ER were also positive for synaptopodin. Also, in primary hippocampal neurons synaptopodin labels the SA and colocalizes with the ER-associated store-operated calcium entry channels STIM1 and Orai-1 (Korkotian et al., 2014). We therefore conclude that the presence of synaptopodin is a suitable marker for the SA. First, we verified the specificity of our synaptopodin antibody using brain sections from synaptopodin deficient mice and established the timeline of synaptopodin expression in rat and mouse hippocampal primary neurons. In agreement with the literature, we found that synaptopodin levels and the degree of spinous localization increased with the maturation of the cultures (Czarnecki et al., 2005; Mundel et al., 1997) Interestingly, when we performed time-lapse imaging of GFP-synaptopodin transfected neurons, we did not observe any processive movements over 7 hours of continued imaging. Varying the time interval between the acquired frames from 10 sec to 10 min we also found no evidence of any active transport of synaptopodin puncta, indicating that under basal conditions, synaptopodin clusters are stably localized and immobile. On the other hand, we observed that new synaptopodin clusters emerged and persisted over extended time periods in dendritic spines. Sometimes this *de novo* formation correlated with ER entry into a spine. It is likely that synaptopodin accumulates in selected spines via association with the actin cytoskeleton from a diffusible pool, or that is locally enriched by specific targeting and translation of synaptopodin mRNA. We speculate that this accumulation is synapse-specific and could depend on the phosphorylation status of synaptopodin. Indeed, in renal podocytes synaptopodin is phosphorylated by PKA and CamKII, this promotes its association with the 14-3-3 protein and protects it from degradation (Faul et al., 2008). Activation of PKA and CamKII pathways in a subset of dendritic spines could lead to selective accumulation of synaptopodin. In our experiments we could not discriminate whether accumulation of synaptopodin precedes or follows ER entering into the spine. Both scenarios are possible and need to be further investigated in the future.

Until now there is a limited number of binding partners known for synaptopodin. Among them are the actin cross-linking proteins α-actinins (Asanuma et al., 2005). Some of the α-actinin isoforms are expressed in neurons and highly enriched in dendritic spines (Hodges et al., 2014; Nakagawa et al., 2004), however, the ubiquitous expression of α-actinins does not explain the selective targeting of synaptopodin to dendritic spines. Taking this into account we decided to search for new binding partners of synaptopodin that could shed some light on the biology of the spine apparatus. Using eukaryotically produced synaptopodin as bait we analyzed the brain-specific interactome of synaptopodin by mass spectrometry. In agreement with our live imaging data, there were no microtubule-based motor proteins or their adaptor proteins found in the synaptopodin complex. On the other hand, we detected a number of unique peptides corresponding to CaMKIIα and CaMKIIβ. Also several 14-3-3 proteins were enriched in the synaptopodin pull down as compared to the control, which suggest that similarly to podocytes, neuronal synaptopodin may be protected from cathepsin L-mediated degradation by association with 14-3-3. The most interesting and the most prominent interaction partners identified in this screen were several F-actin-based myosin motors. Processive myosin V and VI play an important role in a wide range of neuron-specific functions including organelle and mRNA transport and anchoring (Correia et al., 2008; Esteves da Silva et al., 2015). Class V myosins represented by myosin Va and Vb walk towards the fast-growing barbed end of the actin filament and mediate spinous transport of membrane organelles, such as recycling endosomes or tubules of smooth ER (Wagner et al., 2011a; Wang et al., 2008) Moreover, myosin V processivity is regulated by calcium and calmodulin which suggests that myosin function could be controlled by synaptic activity. Myosin VI is the only myosin motor walking in the opposite direction towards the pointed end of the filament. Recruitment of myosin VI to Rab11 recycling endosomes is sufficient to remove them from the synapse (Esteves da Silva et al., 2015). These features make myosins very attractive candidates for synaptic targeting of synaptopodin. Using dominant-negative constructs, myosin VI deficient mice and pharmacological inhibitors of myosin V and VI we tested their effects on synaptopodin localization and motility. We found that interference with myosin V but not myosin VI reduced the number of synaptopodin clusters.

At the moment it is not clear which event is the first step to SA assembly in a selected spine: enrichment of synaptopodin protein, or the entering of an ER protrusion. Since myosin V is required for synaptic targeting of ER in Purkinje neurons, it could be instrumental for initially localizing an ER protrusion to a selected spine, followed by accumulation of synaptopodin and formation of the SA (Figure 7G). Alternatively, is it conceivable that first stabilization of synaptopodin clusters via myosin V and subsequent capture of a transient ER protrusion are necessary for SA assembly. Both mechanisms could co-exist since we found some dendritic protrusions which are positive for the synaptopodin but do not contain ER. Considering that the activity of myosin V is calcium-dependent, in the future it would be interesting to test whether the SA relocates in response to synaptic activity, as this could provide selectivity of synaptic targeting. Together, the results of this study extend our understanding on how synaptopodin and presumably the SA are targeted to the synapses via association with myosin V and actin filaments.

## MATERIALS AND METHODS

### Constructs

To produce the pEGFP-bio-synaptopodin construct, synaptopodin (mouse isoform 3, identifier: Q8CC35-3) was subcloned from a pEGFP-Synaptopodin plasmid (Asanuma et al., 2005) into pEGFP-C1-bio (kind gift from Anna Akhmanova) with SacII and SalI restriction. For cloning of the hSyn1-GFP-synaptopodin construct, GFP-synaptopodin was amplified via PCR, pAAV-hSyn1-mRuby2 was digested with EcoR1 and HindIII to remove mRuby2. GFP-synaptopodin was inserted into the digested plasmid using homologous recombination (Jacobus and Gross, 2015). To produce the MyoVI DN construct, the Syn-GFP-synaptopodin vector was digested with HindIII and NdeI to remove synaptopodin, and the C-terminal domain of mouse myosin VI (bp 3177-3789; NCBI reference sequence: NM_001039546.2) was amplified via PCR and cloned into the backbone (Aschenbrenner et al., 2003).

Purkinje-specific expression plasmids pL7, pL7-mCER, pL7-mRFP-ER-IRES-GFP were described previously (Wagner et al., 2011b, 2011a). Plasmid pL7-mCER-Myo5a-CC-GTD corresponds to pL7-mCER containing a cDNA encoding the C-terminal part of mouse MYO5A (starting at aa residue 1194 of the brain-spliced isoform, transcript variant X6; NCBI reference sequence: XM_006510832.3) inserted in frame at the 3’-end of the mCerulean coding sequence. Similarly, pL7-mCER-Myo5a-GTD carries a cDNA encoding the globular tail domain of mouse MYO5A (starting at aa residue 1415, numbering according to brain-spliced isoform). Plasmid pL7-mCER-LZ was generated by inserting a sequence encoding the leucine zipper of *GCN4* (MKQLEDKVEELLSK NYHLENEVARLKKLVGE) in frame at the 3’-end of the mCerulean coding sequence. To generate pL7-mCER-LZ-Myo5a-GTD, a sequence encoding the globular tail domain of mouse MYO5A (starting at residue 1415, numbering according to brain-spliced isoform) was inserted in frame at the 3’-end of the leucine zipper of pL7-mCER-LZ. Complete list of expression construct used in this study is provided below.

| Backbone | Promoter | Insert | Source |
| --- | --- | --- | --- |
| pAAV | synapsin | mRuby2 | Addgene #99126 (Chan et al., 2017) |
| pAAV | synapsin | synaptopodin-GFP | This study |
| pAAV | synapsin | myosinVI(3433-3705)-DN-GFP | This study |
| pMH4 | synapsin | pDsRed2-ER | Gift from T.G. Oertner |
| pMH4 | synapsin | ER-GFP | (Holbro et al., 2009) |
| pCI | synapsin | tdimer2 | (Campbell et al., 2002) |
| pCI | synapsin | mCerulean | (Wiegert et al., 2018) |
| pCI | synapsin | myosinV-DN-mCerulean | This study |
| pCI | CMV | HA-BirA | (Jaworski et al., 2009) |
| pEGFP | CMV | synaptopodin-GFP-bio | This study |
| pL7 | L7 | empty | (Wagner et al., 2011b) |
| pL7 | L7 | mCerulean | (Wagner et al., 2011b) |
| pL7 | L7 | mRFP-ER-IRES-GFP | (Wagner et al., 2011b) |
| pL7 | L7 | mCER-Myo5a-CC-GTD | This study |
| pL7 | L7 | mCER-Myo5a-GTD | This study |
| pL7 | L7 | mCER-LZ-Myo5a-GTD | This study |

### Antibody

#### Primary antibodies

IF = Immunofluorescence. WB = western blot.

| Antibody | Species | Source | Product # | Dilution |
| --- | --- | --- | --- | --- |
| Synaptopodin | Rabbit | Synaptic Systems | 163002 | IF 1:600 (Fig. 2;4;6;7) |
| Synaptopodin | Mouse | Synaptic Systems | n.a. | IF 1:500 (Fig. S2) |
| Synaptopodin | Rabbit | Sigma | S9442 | IF: 1:1000 (Fig. 1) |
| MAP2 | Mouse | Sigma | M4403 | IF 1:500 (Fig. S2) |
| MAP2 | Rabbit | Abcam | Ab32454 | IF 1:500 (Fig. 2;4;6) |
| homer1 | Mouse | Synaptic Systems | 160011 | IF 1:500 (Fig. 2;6) |
| myosin Va | Rabbit | Sigma | m4812 | WB 1:1000 (Fig. 5) |
| myosin Vb | Rabbit | Santa Cruz | sc-98020 | WB: 1:500 (Fig. 5) |
| myosin Id | Rabbit | Santa Cruz | sc-66982 | WB 1:200 (Fig. 5) |
| myosin VI | Rabbit | Santa Cruz | sc-50461 | WB 1:200 (Fig. 5) |

#### Secondary antibodies

Secondary antibodies:
| Antibody | Conjugation | Source | Product # | Dilution |
| --- | --- | --- | --- | --- |
| anti-rabbit | Alexa Fluor 488 | Life Technologies | A11034 | IF 1:1000 (Fig.1)<br>1:500 (others) |
| anti-rabbit | Alexa Fluor 586 | Life Technologies | A11036 | IF 1:500 (Fig. 2;6;7;S2) |
| anti-mouse | Alexa Fluor 488 | Life Technologies | A11029 | IF 1:500 (Fig. 4) |
| anti-mouse | Alexa Fluor 586 | Life Technologies | A11004 | IF 1:500 (Fig. 2;6;7;S2) |
| Anti-mouse | AbberiorStar 580 | Sigma | 52403 | IF: 1:500 (Fig. 4) |
| anti-rabbit | HRP | Dianova | 111-035-144 | WB 1:20000 (Fig. 5) |

### Pharmacological components

| Reagent | Source | Product number | Final concentration |
| --- | --- | --- | --- |
| MyoVin | Merck | 475984 | 30 μM |
| TIP | Sigma-Aldrich | 19566 | 4 μM |
| phalloidin–Atto647N | Sigma-Aldrich | 65906 | IF: 1:40 |
| Streptavidin-HRP | Pierce | 21130 | WB: 1:10000 |
| DMSO | Carl Roth | 4720.4 | n.a. |

### Animals

Wistar rats Crl:WI(Han) (Charles River) and Wistar Unilever HsdCpb:WU (Envigo) rats were used in this study. Sacrificing of pregnant rats (E18) for primary hippocampal cultures, female P7 rat pups for organotypic slice cultures and adult female rats for biochemistry as well as experiments on mice outlined below were carried out in accordance with the European Communities Council Directive (20110/63/EU) and the Animal Welfare Law of the Federal Republic of Germany (Tierschutzgesetz der Bundersrepublik Deutschland, TierSchG) approved by the local authorities of the city-state Hamburg (Behörde für Gesundheit und Verbraucherschutz, Fachbereich Veterinärwessen, from 21.04.2015) and the animal care committee of the University Medical Center Hamburg-Eppendorf. The mice used in this study were bred and maintained in the animal facility of the Center of Molecular Neurobiology of Hamburg ZMNH, Hamburg, Germany. Synaptopodin knockout mice were described previously (Deller et al., 2003). Perfusions of synaptopodin mice for immunohistochemistry were carried out under license Org 796. *Snell’s waltzer* mice carrying a spontaneously arisen ∼1.0 kb genomic deletion in *Myo6* were obtained from The Jackson Laboratory (B6 x STOCK *Tyr*^*c-ch*^ *Bmp5*^*se*^ *+/+ Myo6*^*sv*^/J; stock no. 000578) and were repeatedly backcrossed to C57BL/6J to obtain *Myo6*^*sv/+*^mice carrying the *Snell’s waltzer* allele but lacking the *Tyr*^*c-ch*^and *Bmp5*^*se*^ alleles (Avraham et al., 1995).

### Perfusion of synaptopodin knockout mouse brains

Adult female mice (11 weeks old for the wild type and 13 weeks old for the knock out) were deeply anesthetized in a gas chamber (20% O^2^ and 80% CO^2^) and sacrificed with 100% CO_2_. Mice were transcardially perfused with phosphate buffered saline followed by fixative (4% PFA + 4% sucrose in PBS, pH 7.4). Brains were removed and post-fixed in the same fixative for a few hours at 4°C. Fixed brains were sectioned using a cryostat (Leica CM3050 S) into 50 μm slices that were kept in cold Phosphate Buffer Saline (PBS). Individual sections were collected with a brush and kept in a 24-well plate in PBS at 4ºC for further immunohistochemistry.

### Organotypic hippocampal cultures, electroporation and 2-photon microscopy

Hippocampal slice cultures were prepared using female rat pups (Wistar) at postnatal day 5–6. The hippocampus was dissected as described (Gee et al., 2017) and cut in 400 µm thick slices using a McIlwain tissue chopper. No antibiotics were used during the preparation or in the culture medium. At DIV 7-9, cultures were transfected with a 1:1.5 mixture of expression vectors encoding tdimer2 (Campbell et al., 2002) and ER-EGFP (Holbro et al., 2009) each driven by the synapsin-1 promoter, using a Helios gene gun (Bio Rad). Imaging of organotypic hippocampal cultures was performed 2-3 weeks after gene transduction. Z-stacks of apical dendrites from CA1 expressing neurons were acquired using a custom-built two-photon imaging setup. It was based on an Olympus BX51WI microscope equipped with a LUMFLN 60×W 1.1 NA objective (Olympus), controlled by the open-source software package ScanImage (Pologruto et al., 2003) written in Matlab (The Mathworks). A pulsed Ti:Sapphire laser (MaiTai DeepSee, Spectra Physics) controlled by electro-optic modulator (350-80, Conoptics) was used to excite tdimer2 and ER-EGFP simultaneously at 980 nm. Emitted photons were collected through objective and oil-immersion condenser (1.4 NA, Olympus) with two pairs of photomultiplier tubes (PMTs, H7422P-40, Hamamatsu). 560 DXCR dichroic mirrors and 525/50 and 607/70 emission filters (Chroma Technology) were used to separate green and red fluorescence. Excitation light was blocked by short-pass filters (ET700SP-2P, Chroma). Slices were submerged in artificial cerebrospinal fluid (ACSF) containing (in mM): 127 NaCl, 2.5 KCl, 2 CaCl_2_, 1 MgCl_2_, 25 NaHCO_3_, 1.25 NaH_2_PO_4_, 25 D-glucose (pH 7.4, ∼308 mOsm, saturated with 95% O_2_/ 5% CO_2_) at RT. After 2P live imaging, slices were fixed overnight at 4°C in a 2% (w/v) PFA/sucrose mixture and used for post hoc staining with synaptopodin antibody.

### Immunohistochemistry and imaging of organotypic slices

For immunostaining, the slices were washed 3 x in PBS, followed by overnight permeabilization with 1% Triton-X-100 at room temperature (RT). In between each of the following steps using PBS-based solutions the slices were washed in PBS. For quenching, slices were submerged in Glycine (50 mM) for 1 h. Afterwards, ImageIT Fx enhancer (Thermo Fisher) was used to avoid unspecific labelling. Rat hippocampal slices were incubated with rabbit primary antibody (1:1000) against rat synaptopodin (S9442, from Sigma) followed for 24 h at 4°C. Next, the slices were incubated with secondary antibody Alexa488 goat-anti-rabbit (1:1000) for 12 h at 4°C. Finally, the slices were washed and mounted using the Prolong Antifade Kit (Thermo Fisher). Images were taken using an Olympus FV300 confocal microscope equipped with an UPLSAPO 60× 1.35 NA objective and analyzed with Fiji (Schindelin et al., 2012). For immunostaining of synaptopodin +/+ and -/- brain sections were fixed 5 minutes in 4% Roti-Histofix (Carl Roth), 4% Sucrose and after washed 3 times in PBS followed by permeabilization with 1% Triton-X-100 at room temperature for 1 hour. After 3 x washing in PBS, slices were incubated in blocking buffer, BB (2% BSA, 0,5% Triton in PBDS) for 1.5h at RT. Incubation with primary antibodies in BB was performed overnight at 4º C. Brain sections were again washed 3 times in PBS and after incubated with the corresponding secondary antibody in PBS for 1h at RT. Before mounting in mowiol, slices were shortly incubated for 10 min with DAPI.

### Primary neuronal culture and transfections

*Primary hippocampal rat cultures* were essentially prepared as described previously (Kapitein et al., 2010). In brief, hippocampi were dissected from E18 embryos, treated with trypsin (0.25 %, Thermo Fisher Scientific) for 10 minutes at 37°C, physically dissociated by pipetting through a syringe, and plated on 18 mm glass coverslips coated with poly-L-lysine (Sigma-Aldrich) at a density of 40000-60000 cells/ml and coverslip in DMEM (Gibco) with 10 % fetal calf serum (Gibco) and Penicillin/streptomycin (Thermo Fisher Scientific). After 1 h, the medium was replaced by BrainPhys neuronal medium supplemented with SM1 (Stem Cell kit) and 0.5 mM glutamine (Thermo Fisher Scientific) without antibiotics. Cells were then kept in an incubator at 37 °C, 5 % CO_2_and 95 % humidity. Primary hippocampal rat cultures were transfected with lipofectamine 2000 (Thermo Fisher Scientific) according to the manufacturer’s introductions and as described previously described (Kapitein et al., 2010).

*Primary hippocampal mouse cultures* from *Myo6*^*sv/sv*^ and *Myo6*^*+/+*^ mice were prepared as described (Spilker et al., 2016). Briefly, hippocampi were dissected from male and female P0 *Myo6*^*sv/sv*^ and *Myo6*^*+/+*^ mice and cells were dissociated after 10 min treatment with trypsin at 37 °C. Neurons were plated on glass dishes coated with poly-L-lysine (Sigma-Aldrich) at a density of 30.000 cells per well /12 well plate) in DMEM medium (Gibco) supplemented with 8 % FCS, 1% penicillin/streptomycin. Following attachment, mouse cultures were kept in Neurobasal medium (Gibco) supplemented with 2 mM Glutamine, 1 % penicillin/streptomycin and 1X B27 supplement (Gibco), at 37 °C, 5 % CO_2_and 95 % humidity.

*Dissociated cerebellar cultures* containing Purkinje cells were prepared and transfected as described in detail (Wagner et al., 2011b).

### Production of AAV and infection of primary hippocampal neurons

Adeno-associated viruses were produced at the Vector Facility of the University Medical Center Hamburg-Eppendorf (UKE). pAAV-hSyn1-mRuby2 (Addgene 99126) and pAAV-hSyn1-GFP-synaptopodin (this study) were respectively packaged by pE2/rh10 and p5E/9 (Julie C. Johnston, University of Pennsylvania, USA) and pHelper (CellBiolabs). For infection with AAV, the viruses were added directly into the culture medium at final concentrations between 10^11^-10^9^ vg/ml. pSyn-GFP-Synaptopodin AAV9 was added to the culture at DIV10, mRuby2 AAVrh10 was added on DIV14, and cells were imaged on DIV17.

### Immunocytochemistry

Cells were fixed in 4 % Roti-Histofix (Carl Roth), 4 % Sucrose in PBS for 10 min at RT and washed three times with PBS, before they were permeabilized in 0.2 % Triton X-100 in PBS for 10 min. After 3 x washing in PBS, coverslips were incubated in blocking buffer (BB, 10% horse serum and 0.1% Triton in PBS) for 1 h at RT. Incubation with primary antibodies was performed in BB at 4 °C overnight. After 3 x washing in PBS, cells were incubated with corresponding secondary antibodies in BB for 1 h at RT. If the staining included phalloidin, an additional step was added where the coverslips were incubated with phalloidin-647N (1:40) overnight at 4 °C. Finally, the coverslips were washed 5 x in PBS and mounted in mowiol.

### Confocal microscopy: fixed and live cell imaging

Z-stack images of fixed primary hippocampal neurons were acquired on a Leica TCS SP8 confocal microscope using 63.0 x 1.40 oil objective using 488 nm, 568 nm and 633 nm excitation lasers. The pixel size was set to 90 nm and z-steps varied between 250-350 nm. For the shown representative confocal images, a Gaussian filter (radius 0.5 px) was applied in ImageJ to reduce the visible background noise. Live Purkinje cells were imaged at 37ºC on a Zeiss LSM 510 confocal microscope equipped with a 100 x 1.40 oil objective exactly as described previously (Wagner et al., 2011a, 2011b).

### Analysis and quantification of synaptopodin and homer1 puncta

The quantification and distribution of synaptopodin and homer1 puncta in primary hippocampal neurons was analyzed from confocal z-stack images using Fiji/ImageJ. Z-stacks were projected into one plane using the Z projection function (Max Intensity) and decoded for blind analysis by an independent researcher. 1 – 4 dendritic segments of approximately 40-60 µm length were selected per cell and used for quantification of the number of synaptopodin and homer1 puncta. Data were averaged per cell. “Spines” were defined as protrusions from the main dendrite labeled by phalloidin or MAP2 and were positive for post-synaptic marker homer1. All other puncta that appeared to be localized in the dendritic shaft were defined as “inside dendrite”. A small percentage of synaptopodin puncta appeared to be localized outside of the dendritic shaft in spine-like protrusions (or filopodia) that did not contain homer1 staining. Those were defined as “inside protrusion”. For images that did not contain homer1 staining, synaptopodin puncta were defined as “inside” or “outside” the dendritic shaft, i.e. the “outside” fraction contains both spinous and filopodia fractions.

### STED Imaging

Confocal and STED images of phalloidin (Alexa647), MAP2 (Alexa488) and synaptopodin (Abberior Star 580) were acquired on a Leica TCS SP8-3X gated STED microscope equipped with a pulsed 775 nm depletion laser and a pulsed white light laser (WLL) for excitation. For acquiring images, the Leica objective HC APO CS2 100x/1.40 oil was used. Fluorescence of the respective channel was excited by the WLL at 650 nm (STED and confocal), 488 nm (confocal mode) and 561 nm (STED and confocal), respectively. For STED imaging, emission was acquired between 660-730 nm for Alexa-647 and 580-620 nm for Abberior Star 580. The detector time gates for both STED channels were set from 0.5-1 ns to 6 ns. Both dyes were depleted with 775 nm. Images were taken as single plane of 1024×1024 pixels and digital zoom factor 5.

### Wide field and TIRF microscopy: live cell imaging

Live cell wide-field and TIRF imaging for GFP-synaptopodin motility analysis was performed on a Nikon Eclipse Ti-E controlled by VisiView software (VisitronSystems). The microscope was equipped with 488 nm, 561 nm, and 640 nm excitation laser lines via a multi-mode fiber. Oblique and TIRF illuminations were achieved with an ILAS2 TIRF system. The angle of the excitation light was adjusted manually to achieve optimal signal/noise ratio. Samples were imaged with a 100x TIRF objective (Nikon, ApoTIRF 100x/1.49 oil). Emission light was captured through a quad-band filter (Chroma, 405/488/561/640) followed by a filter wheel with filters for GFP (Chroma, 525/50m), RFP (Chroma, 595/50m), and Cy5 (Chroma, 700/75m). Multi-channel images were acquired sequentially with an Orca flash 4.0LT CMOS camera (Hamamatsu).

### Synaptopodin motility analysis

Motility analysis was done on a time lapse series of DIV16 primary hippocampal neurons expressing GFP-synaptopodin and mRuby2 (AAV infection). ∼10 cells were selected, their position was saved, one image was acquired in the red channel (mRuby2) and a time lapse was acquired in the green channel (GFP-synaptopodin) over a 5 minute period with 10 second intervals. The medium was then supplemented with either 30 µM MyoVin or 4 µM TIP. Cells were incubated for ∼45 minutes with the inhibitor before the described imaging regime was repeated. For the analysis, cells that were still healthy after the incubation period (mRuby2 channel; no blebbing or fragmentation visible) were selected. Motility of synaptopodin-puncta within the same cell before and after treatment was assessed using an ImageJ script as described in (Verkuyl and Matus, 2006). In brief, the minimum – maximum display range was set to 0 – 65535 for all image stacks. Then a motility plot was generated from the time lapse stack by summing up the differences in grayscale values for each pixel from frame to frame as described by Verkuyl & Matus. The overall intensity of such generated motility plots was measured for the whole image (integrated density measurement), representing the overall motility observed in the time lapse stack.

### Culturing and transfection of HEK293 cells

HEK293T cells (from Mikhaylova et al., 2018) were maintained in full medium consisting of Dulbecco’s modified Eagle’s medium (DMEM; GIBCO, Thermo Fisher) supplemented with 10 % fetal calf serum (FCS), 1 x penicillin/streptomycin and 2 mM glutamine at 37 °C, 5 % CO_2_ and 95 % humidity. For the expression of biotinylated proteins, HEK293T cells were grown in full medium made with a 50 % DMEM, 50 % Ham’s F-10 Nutrient Mix (GIBCO, Thermo Fisher) mixture. Transfection was done using MaxPEI 25K (Polysciences) in a 3:1 MaxPEI:DNA ratio.

### Pull-down for western blot and mass spec analysis

HEK293 cells were co-transfected with HA-BirA and either Synaptopodin-pEGFP-bio or pEGFP-bio (control vector) and harvested after 18 h of expression. The cells were lysed in extraction buffer (20 mM Tris pH 8, 150 mM NaCl, 1% Triton-X-100, 5 mM MgCl_2_, complete protease inhibitor cocktail (Roche)), kept on ice for 30 min and centrifuged for 15 min at 14000 g. Magnetic Streptavidin M-280 Dynabeads (Invitrogen) were washed 3 x in washing buffer (20 mM Tris pH 8, 150 mM KCl, 0.1 % Triton-X 100), blocked in 3 % chicken egg albumin (Sigma) for 40 min at RT, and again washed 3 x in washing buffer. The cleared cell lysate was added to the blocked beads and incubated at 4 °C on a rotator overnight. After the incubation period, the beads were washed 2 x with low salt washing buffer (100 mM KCl), 2 x in high salt washing buffer (500 mM KCl) and again 2 x in low salt washing buffer. For preparation of rat brain extract, 9 ml lysis buffer (50mM Tris HCl pH 7.4, 150 mM NaCl, 0.1 % SDS, 0.2 % NP-40, complete protease inhibitor cocktail (Roche)) were added per 1 g of tissue weight, and the tissue was lysed using a Dounce homogenizer. The lysate was cleared first for 15 min at 1000 g, and the supernatant was again centrifuged for 20 min at 15000 g to obtain the final lysate. The washed beads were then combined with 1 ml of the cleared rat brain lysate and incubated for 1 h at 4 °C on a rotator. Finally the beads were washed 5 x in washing buffer and resuspended in Bolt LDS sample buffer (Invitrogen) for subsequent SDS-PAGE. For mass-spectrometric analysis, the samples were separated on a commercial Bolt Bis-Tris Plus Gel (Invitrogen) and the intact gel was sent to Erasmus MC Proteomics Center, Rotterdam, for mass spectrometry analysis. For Western blot analysis, the samples were loaded on 4 % - 20 % acrylamide gradient gels and blotted on PVDF membranes. After blocking in 5 % skim milk in Tris-buffered saline (TBS, 20 mM Tris pH 7.4, 150 mM NaCl, 0.1 % Tween-20), membranes were incubated with primary antibodies diluted in TBS-A (TBS pH 7.4, 0.02 % sodium-azide) overnight at 4 °C. Corresponding HRP-conjungated secondary antibodies and HRP-Strepdavidin were applied for 1.5 h at RT in 5 % skim milk in TBS.

### Mass spectrometry analysis

SDS-PAGE gel lanes were cut into 1-mm slices using an automatic gel slicer. Per sample 8 to 9 slices were combined and subjected to in-gel reduction with dithiothreitol, alkylation with iodoacetamide and digestion with trypsin (Thermo Fisher Scientific, TPCK treated), essentially as described by (Wilm et al., 1996). Nanoflow LCMS/MS was performed on an EASY-nLC 1000 Liquid Chromatograph (Thermo Fisher Scientific) coupled to an Orbitrap Fusion™ Tribrid™ Mass Spectrometer (Thermo Fisher Scientific) operating in positive mode and equipped with a nanospray source. Peptide mixtures were trapped on a nanoACQUITY UPLC C18 column (100Å, 5 µm, 180 µm X 20 mm, Waters). Peptide separation was performed on a ReproSil C18 reversed phase column (Dr Maisch GmbH; 20 cm × 100 µm, packed in-house) using a linear gradient from 0 to 80% B (A = 0.1 % formic acid; B = 80% (v/v) acetonitrile, 0.1 % formic acid) in 60 min and at a constant flow rate of 500 nl/min. Mass spectra were acquired in continuum mode; fragmentation of the peptides was performed in data-dependent mode. Peak lists were automatically created from raw data files using the Mascot Distiller software (version 2.1; MatrixScience). The Mascot search algorithm (version 2.2, MatrixScience) was used for searching against a UniProt canonical database (release 2016_07, taxonomies Homo sapiens and Rattus norvegicus combined). The peptide tolerance was typically set to 10 ppm and the fragment ion tolerance to 0.8 Da. A maximum number of 2 missed cleavages by trypsin were allowed and carbamidomethylated cysteine and oxidized methionine were set as fixed and variable modifications, respectively. Quantitative analysis was done with MaxQuant software (version 1.5.4.1) using similar or standard settings and using the UniProt rat and human isoforms databases (both releases 2016_07).

### Network analysis of potential synaptopodin-interactors

We selected 50 proteins with the highest unique peptide count from the mass spectrometry analysis of synaptopodin interactors for the network analysis using the “Multiple Proteins” function of the online STRING database (http://string-db.org; Szklarczyk et al., 2015). The parameters changed from the default settings were as follows: Network edges: “Confidence” (line thickness indicates the strength of data support). Active interaction sources: “Experiments” and “Databases”. Selected network statistics / Functional enrichments: Biological Process (GO)

### Statistical analysis

Statistical analysis was performed in Prism 6.05 (GraphPad, LA Jolla, CA, USA) or in Statistica 13 (Dell Inc). Detailed information about the type of test used, significance levels, n numbers and biological replicates are provided in the figure legends. All averaged data are shown as mean ± SEM. The analysis of the *Myo6*^*sv/sv*^ and *Myo6*^*+/+*^ mice data was done blindly.

## ACKNOWLEGEMENTS

The authors would like to thank Dr. Antonio Virgilio Failla from the UKE Microscopy Imaging Facility and Oliver Kobler from the Combinatorial Neuroimaging Core Facility at LIN Magdeburg for access to STED microscope and Dr. Ingke Braren from the Vector Facility of the UKE for generation of adeno-associated viruses. pAAV-hSyn1-mRuby2 was a gift from Viviana Gradinaru (Addgene plasmid # 99126; http://n2t.net/addgene:99126; RRID:Addgene_99126).

## DATA AVAILABILITY

Mass spectrometry results (Table S1) will be uploaded to *Dryad*.

## REFERENCES

Asanuma, K., Kim, K., Oh, J., Giardino, L., Chabanis, S., Faul, C., et al. (2005). Synaptopodin regulates the actin-bundling activity of α-actinin in an isoform-specific manner. J. Clin. Invest. 115, 1188–1198. doi:10.1172/JCI200523371.

Aschenbrenner, L., Lee, T., and Hasson, T. (2003). Myo6 Facilitates the Translocation of Endocytic Vesicles from Cell Peripheries. Mol. Biol. Cell 14, 2728–2743.

Avraham, K. B., Hasson, T., Steel, K. P., Kingsley, D. M., Russel, L. B., Mooseker, M. S., et al. (1995). The mouse Snell’s waltzer deafness gene encodes an unconventional myosin required for structural integrity of inner ear hair cells. Nature 11, 369–375. doi:10.1038/ng1295-369.

Ayloo, S., Guedes-Dias, P., Ghiretti, A. E., and Holzbaur, E. L. F. (2017). Dynein efficiently navigates the dendritic cytoskeleton to drive the retrograde trafficking of BDNF/TrkB signaling endosomes. Mol. Biol. Cell 28, 2543–2554. doi:10.1091/mbc.E17-01-0068.

Balasanyan, V., and Arnold, D. B. (2014). Actin and Myosin-dependent localization of mRNA to dendrites. PLoS One 9, e92349. doi:10.1371/journal.pone.0092349.

Bas Orth, C., Schultz, C., Müller, C. M., Frotscher, M., and Deller, T. (2007). Loss of the Cisternal Organelle in the Axon Initial Segment of Cortical Neurons in Synaptopodin-Deficient Mice. J. Comp. Neurol. 504, 441–449.

Campbell, R. E., Tour, O., Palmer, A. E., Steinbach, P. A., Baird, G. S., Zacharias, D. A., et al.. (2002). A monomeric red fluorescent protein. Proc. Natl. Acad. Sci. 99, 7877–7882. doi:10.1073/pnas.082243699.

Chalovich, J. M., and Schroeter, M. M. (2010). Synaptopodin family of natively unfolded, actin binding proteins: Physical properties and potential biological functions. Biophys. Rev. 2, 181–189. doi:10.1007/s12551-010-0040-5.

Chan, K. Y., Jang, M. J., Yoo, B. B., Greenbaum, A., Ravi, N., Wu, W.-L., et al.. (2017). Engineered AAVs for efficient noninvasive gene delivery to the central and peripheral nervous systems. Nat. Neurosci. 20, 1172–1179. doi:10.1038/nn.4593.

Correia, S. S., Bassani, S., Brown, T. C., Lisé, M.-F., Backos, D. S., El-Husseini, A., et al.. (2008). Motor protein-dependent transport of AMPA receptors into spines during long-term potentiation. Nat. Neurosci. 11, 457–66. doi:10.1038/nn2063.

Czarnecki, K., Haas, C. A., Bas Orth, C., Deller, T., and Frotscher, M. (2005). Postnatal development of synaptopodin expression in the rodent hippocampus. J. Comp. Neurol. 490, 133–144. doi:10.1002/cne.20651.

Deller, T., Bas Orth, C., Del Turco, D., Vlachos, A., Burbach, G. J., Drakew, A., et al.. (2007). A role for synaptopodin and the spine apparatus in hippocampal synaptic plasticity. Ann. Anat. 189, 5–16. doi:10.1016/j.aanat.2006.06.013.

Deller, T., Korte, M., Chabanis, S., Drakew, A., Schwegler, H., Stefani, G. G., et al.. (2003). Synaptopodin-deficient mice lack a spine apparatus and show deficits in synaptic plasticity. Proc. Natl. Acad. Sci. U. S. A. 100, 10494–9. doi:10.1073/pnas.1832384100.

Deller, T., Merten, T., Roth, S. U., Mundel, P., and Frotscher, M. (2000). Actin-associated protein synaptopodin in the rat hippocampal formation: Localization in the spine neck and close association with the spine apparatus of principal neurons. J. Comp. Neurol. 418, 164–181. doi:10.1002/(SICI)1096-9861(20000306)418:2<164:AID-CNE4>3.0.CO;2-0.

Esteves da Silva, M., Adrian, M., Schätzle, P., Lipka, J., Watanabe, T., Cho, S., et al.. (2015). Positioning of AMPA Receptor-Containing Endosomes Regulates Synapse Architecture. Cell Rep. 13, 933–943. doi:10.1016/j.celrep.2015.09.062.

Faul, C., Donnelly, M., Merscher-Gomez, S., Chang, Y. H., Franz, S., Delfgaauw, J., et al.. (2008). The actin cytoskeleton of kidney podocytes is a direct target of the antiproteinuric effect of cyclosporine A. Nat. Med. 14, 931–938. doi:10.1038/nm.1857.

Gee, C. E., Ohmert, I., Wiegert, J. S., and Oertner, T. G. (2017). Preparation of Slice Cultures from Rodent Hippocampus. Cold Spring Harb. Protoc. 2017, pdb.prot094888. doi:10.1101/pdb.prot094888.

Hanus, C., and Ehlers, M. D. (2016). Specialization of biosynthetic membrane trafficking for neuronal form and function. Curr. Opin. Neurobiol. 39, 8–16. doi:10.1016/j.conb.2016.03.004.

Hanus, C., Kochen, L., Tom Dieck, S., Racine, V., Sibarita, J. B., Schuman, E. M., et al.. (2014). Synaptic Control of Secretory Trafficking in Dendrites. Cell Rep. 7, 1771–1778. doi:10.1016/j.celrep.2014.05.028.

Hodges, J. L., Vilchez, S. M., Asmussen, H., Whitmore, L. A., and Horwitz, A. R. (2014). α-Actinin-2 mediates spine morphology and assembly of the post-synaptic density in hippocampal neurons. PLoS One 9. doi:10.1371/journal.pone.0101770.

Holbro, N., Grunditz, A., and Oertner, T. G. (2009). Differential distribution of endoplasmic reticulum controls metabotropic signaling and plasticity at hippocampal synapses. Proc. Natl. Acad. Sci. U. S. A. 106, 15055–60. doi:10.1073/pnas.0905110106.

Jacobus, A. P., and Gross, J. (2015). Optimal cloning of PCR fragments by homologous recombination in Escherichia coli. PLoS One 10, 1–17. doi:10.1371/journal.pone.0119221.

Jaworski, J., Kapitein, L. C., Gouveia, S. M., Dortland, B. R., Wulf, P. S., Grigoriev, I., et al.. (2009). Dynamic Microtubules Regulate Dendritic Spine Morphology and Synaptic Plasticity. Neuron 61, 85–100. doi:10.1016/j.neuron.2008.11.013.

Jones, J. M., Huang, J. D., Mermall, V., Hamilton, B. A., Mooseker, M. S., Escayg, A., et al.. (2000). The mouse neurological mutant flailer expresses a novel hybrid gene derived by exon shuffling between Gnb5 and Myo5a. Hum Mol Genet 9, 821–828. doi:10.4018/IJMCMC.2016100103.

Kapitein, L. C., Schlager, M. A., Kuijpers, M., Wulf, P. S., van Spronsen, M., MacKintosh, F. C., et al.. (2010). Mixed Microtubules Steer Dynein-Driven Cargo Transport into Dendrites. Curr. Biol. 20, 290–299. doi:10.1016/j.cub.2009.12.052.

Korkotian, E., Frotscher, M., and Segal, M. (2014). Synaptopodin Regulates Spine Plasticity: Mediation by Calcium Stores. J. Neurosci. 34, 11641–11651. doi:10.1523/JNEUROSCI.0381-14.2014.

Kremerskothen, J., Plaas, C., Kindler, S., Frotscher, M., and Barnekow, A. (2005). Synaptopodin, a molecule involved in the formation of the dendritic spine apparatus, is a dual actin/??-actinin binding protein. J. Neurochem. 92, 597–606. doi:10.1111/j.1471-4159.2004.02888.x.

Kügler, S., Kilic, E., and Bähr, M. (2003). Human synapsin 1 gene promoter confers highly neuron-specific long-term transgene expression from an adenoviral vector in the adult rat brain depending on the transduced area. Gene Ther. 10, 337–347. doi:10.1038/sj.gt.3301905.

Matt, L., Kim, K., Hergarden, A. C., Patriarchi, T., Malik, Z. A., Park, D. K., et al.. (2018). α-Actinin Anchors PSD-95 at Postsynaptic Sites. Neuron 97, 1094–1109.e9. doi:10.1016/j.neuron.2018.01.036.

Mikhaylova, M., Bär, J., van Bommel, B., Schätzle, P., YuanXiang, P. A., Raman, R., et al.. (2018). Caldendrin Directly Couples Postsynaptic Calcium Signals to Actin Remodeling in Dendritic Spines. Neuron 97, 1110–1125.e14. doi:10.1016/j.neuron.2018.01.046.

Mikhaylova, M., Bera, S., Kobler, O., Frischknecht, R., and Kreutz, M. R. (2016). A Dendritic Golgi Satellite between ERGIC and Retromer. Cell Rep. 14, 189–199. doi:10.1016/j.celrep.2015.12.024.

Mundel, P., Heid, H. W., Mundel, T. M., Krüger, M., Reiser, J., and Kriz, W. (1997). Synaptopodin: An actin-associated protein in telencephalic dendrites and renal podocytes. J. Cell Biol. 139, 193–204. doi:10.1083/jcb.139.1.193.

Nakagawa, T., Engler, J. A., and Sheng, M. (2004). The dynamic turnover and functional roles of α-actinin in dendritic spines. Neuropharmacology 47, 734–745. doi:10.1016/j.neuropharm.2004.07.022.

Osterweil, E., Wells, D. G., and Mooseker, M. S. (2005). A role for myosin VI in postsynaptic structure and glutamate receptor endocytosis. J. Cell Biol. 168, 329–38. doi:10.1083/jcb.200410091.

Pologruto, T. A., Sabatini, B. L., and Svoboda, K. (2003). ScanImage: flexible software for operating laser scanning microscopes. Biomed. Eng. Online 2, 13. doi:10.1186/1475-925X-2-13.

Sánchez-Ponce, D., DeFelipe, J., Garrido, J. J., and Muñoz, A. (2011). In vitro maturation of the cisternal organelle in the hippocampal neuron’s axon initial segment. Mol. Cell. Neurosci. 48, 104–116. doi:10.1016/j.mcn.2011.06.010.

Schindelin, J., Arganda-Carreras, I., Frise, E., Kaynig, V., Longair, M., Pietzsch, T., et al.. (2012). Fiji: an open-source platform for biological-image analysis. Nat. Methods 9, 676–682. doi:10.1038/nmeth.2019.

Spilker, C., Nullmeier, S., Grochowska, K. M., Schumacher, A., Butnaru, I., Macharadze, T., et al.. (2016). A Jacob/Nsmf Gene Knockout Results in Hippocampal Dysplasia and Impaired BDNF Signaling in Dendritogenesis. PLoS Genet. 12, 1–32. doi:10.1371/journal.pgen.1005907.

Szklarczyk, D., Franceschini, A., Wyder, S., Forslund, K., Heller, D., Huerta-Cepas, J., et al.. (2015). STRING v10: Protein-protein interaction networks, integrated over the tree of life. Nucleic Acids Res. 43, D447–D452. doi:10.1093/nar/gku1003.

Toresson, H., and Grant, S. G. N. (2005). Dynamic distribution of endoplasmic reticulum in hippocampal neuron dendritic spines. Eur. J. Neurosci. 22, 1793–1798. doi:10.1111/j.1460-9568.2005.04342.x.

Valenzuela, J., and Perez, F. (2015). Diversifying the secretory routes in neurons. Front. Neurosci. 9. doi:10.3389/fnins.2015.00358.

van Spronsen, M., Mikhaylova, M., Lipka, J., Schlager, M. A., van den Heuvel, D. J., Kuijpers, M., et al.. (2013). TRAK/Milton Motor-Adaptor Proteins Steer Mitochondrial Trafficking to Axons and Dendrites. Neuron 77, 485–502. doi:10.1016/j.neuron.2012.11.027.

Verbich, D., Becker, D., Vlachos, A., Mundel, P., Deller, T., and McKinney, R. A. (2016). Rewiring neuronal microcircuits of the brain via spine head protrusions--a role for synaptopodin and intracellular calcium stores. Acta Neuropathol. Commun. 4, 38. doi:10.1186/s40478-016-0311-x.

Verkuyl, J. M., and Matus, A. (2006). Time-lapse imaging of dendritic spines in vitro. Nat. Protoc. 1, 2399–2405. doi:10.1038/nprot.2006.357.

Vlachos, A., Ikenberg, B., Lenz, M., Becker, D., Reifenberg, K., Bas-Orth, C., et al.. (2013). Synaptopodin regulates denervation-induced homeostatic synaptic plasticity. Proc. Natl. Acad. Sci. 110, 8242–8247. doi:10.1073/pnas.1213677110.

Vlachos, A., Korkotian, E., Schonfeld, E., Copanaki, E., Deller, T., and Segal, M. (2009). Synaptopodin regulates plasticity of dendritic spines in hippocampal neurons. J. Neurosci. 29, 1017–1033. doi:10.1523/JNEUROSCI.5528-08.2009.

Wagner, W., Brenowitz, S. D., and Hammer, J. A. I. (2011a). Myosin-Va Transports the Endoplasmic Reticulum into the Dendritic Spines of Purkinje Neurons. Nat. Cell Biol. 1, 40–48. doi:10.1038/ncb2132.

Wagner, W., McCroskery, S., and Hammer, J. A. (2011b). An Efficient Method for the Long-Term and Specific Expression of Exogenous cDNAs in Cultured Purkinje Neurons. J. Neurosci. Methods 200, 95–105. doi:10.1016/j.jneumeth.2011.06.006.

Wang, L., Dumoulin, A., Renner, M., Triller, A., and Specht, C. G. (2016). The Role of Synaptopodin in Membrane Protein Diffusion in the Dendritic Spine Neck. PLoS One 11, e0148310. doi:10.1371/journal.pone.0148310.

Wang, Z., Edwards, J. G., Riley, N., Provance, D. W., Karcher, R., Li, X., et al.. (2008). Myosin Vb Mobilizes Recycling Endosomes and AMPA Receptors for Postsynaptic Plasticity. Cell 135, 535–548. doi:10.1016/j.cell.2008.09.057.

Wiegert, J. S., Pulin, M., Gee, C. E., and Oertner, T. G. (2018). The fate of hippocampal synapses depends on the sequence of plasticity-inducing events. Elife 7, 1–18. doi:10.7554/eLife.39151.

Wilm, M., Shevchenko, A., Houthaeve, T., Breit, S., Schweigerer, L., Fotsis, T., et al.. (1996). Femtomole sequencing of proteins from polyacrylamide gels by nano-electrospray mass spectrometry. Nature 379, 466–469.

Wu, X., Wang, F., Rao, K., Sellers, J. R., and III, J. A. H. (2002). Rab27a Is an Essential Component of Melanosome Receptor for Myosin Va. Mol. Biol. Cell 13, 2170–2179. doi:10.1091/mbc.01.

Yamazaki, M., Matsuo, R., Fukazawa, Y., Ozawa, F., and Inokuchi, K. (2001). Regulated expression of an actin-associated protein, synaptopodin, during long-term potentiation. J. Neurochem. 79, 192–199. doi:10.1046/j.1471-4159.2001.00552.x.

Zhang, X., Pöschel, B., Faul, C., Upreti, C., Stanton, P. K., and Mundel, P. (2013). Essential role for synaptopodin in dendritic spine plasticity of the developing hippocampus. J. Neurosci. 33, 12510–8. doi:10.1523/JNEUROSCI.2983-12.2013.

